# Intellectual disability-causing mutations in KIF11 impair microtubule dynamics and dendritic arborization

**DOI:** 10.1101/2024.10.02.615913

**Authors:** Jenna L. Wingfield, Lukas Niese, Rahul Grover, Stefan Diez, Sathyanarayanan V Puthanveettil

**Author notes:** Corresponding author: Sathyanarayanan V Puthanveettil.

## Abstract

Precise control of axonal and dendritic architecture is vital for proper brain function, with microtubule (MT) dynamics playing a central role in this process. Here, we uncover a previously unrecognized function of the molecular motor protein KIF11, which acts as a MT dynamics rheostat in hippocampal neurons to modulate dendritic branching. Known for its role in mitotic spindle bipolarity, KIF11 is also implicated in Microcephaly with or without chorioretinopathy, lymphedema, or intellectual disabilities (MCLID). However, the specific neuronal functions of KIF11 and the impact of its mutations in MCLID have remained largely unexplored. Our studies, using quantitative imaging of MT dynamics following KIF11 inhibition, indicate that KIF11 preferentially binds to parallel MTs in mature neurons. This binding is associated with a marked increase in minus-end-out MT dynamics in both axons and dendrites upon KIF11 loss of function, coupled with enhanced MT flux and extended growth in tertiary dendrites. These changes suggest a novel role for KIF11 in orchestrating dendritic branching. Moreover, introducing MCLID-associated KIF11 mutations, KIF11^Y82F^, and KIF11^ΔCterm^, which cause minor microcephaly but severe intellectual disabilities, leads to significantly reduced MT dynamics and impaired dendritic arborization. In a microtubule sliding assay, KIF11^Y82F^ significantly reduced KIF11 velocity while KIF11^ΔCterm^ increased it. Temporal inhibition of KIF11 using a photo-inhibitable KIF11, show increased MT dynamics and dendritic growth, while activation results in kinked and twisted branches. Together, these data reveal that KIF11 is MT dynamics rheostat and regulator of dendritic arborization in mature neurons and provide new insights into the molecular mechanisms driving MCLID.

## INTRODUCTION

Neurons, among the most asymmetric and largest cell types, rely on precise cytoskeletal control and directed long-distance transport for intercellular communication and proper brain wiring. Several studies, including our own, have demonstrated that kinesin superfamily (KIFs) molecular motor proteins are key modulators of both intra- and intercellular communication (Aiken and Holzbaur, 2021; Alsabban et al., 2020; Fan and Lai, 2022; Joseph et al., 2021b; Morikawa et al., 2018; Muhia et al., 2016; Puthanveettil et al., 2008; Swarnkar et al., 2021; Swarnkar et al., 2018; Zhou et al., 2013). These proteins play crucial roles in maintaining neuronal health, as well as in brain development and function (Hirokawa et al., 2010). Most of the KIFs studied in neurons, particularly those involved in learning and memory, are kinesins directly responsible for the long-distance microtubule (MT) dependent transport of molecular and organelle cargos (Guedes-Dias and Holzbaur, 2019; Morikawa et al., 2018; Wang and Xu, 2015). However, emerging research indicates that certain KIFs, traditionally studied in the context of cell division, are also expressed and functional in mature neurons (Joseph et al., 2021a; Silverman et al., 2010). However, the mechanisms underlying their roles in these post-mitotic cells remain poorly understood.

Elucidating the functions and mechanisms of mitotic KIFs in post-mitotic neurons is expected to uncover novel pathways that govern KIFs and neural circuit function. This line of research holds promise for advancing our understanding of disease mechanisms and could open new avenues for therapeutic development.

To address this significant knowledge gap, we investigated the mechanisms and functions of the mitotic KIF, KIF11, in hippocampal neurons. Notably, mutations in the mitotic motor KIF11 (kinesin-5/EG5) have been linked to microcephaly with or without chorioretinopathy, lymphedema, or intellectual disabilities (MCLID) (Balikova et al., 2016; Jones et al., 2014; Malvezzi et al., 2018). MCLID is a rare autosomal dominant disorder characterized by these phenotypic features (Güneş et al., 2018; Jones et al., 2014; Schlögel et al., 2015). Patients with MCLID are heterozygous, and disease-relevant KIF11 mutations, predicted to be loss-of-function, occur throughout the KIF11 gene. However, these predictions have not yet been experimentally confirmed. In rodent models, homozygous KIF11 knockout (KO) mice die before the blastocyst stage, whereas heterozygous mice develop normally, suggesting that KIF11 haploinsufficiency alone is unlikely to account for the observed disease phenotypes (Chauvière et al., 2008; Wang et al., 2020). Consequently, the mechanism by which KIF11 mutations lead to intellectual disabilities remains elusive.

The pathomechanism of MCLID has largely been attributed to KIF11’s role in mitosis. KIF11 is a homotetrameric kinesin that generates force on microtubules (MTs) by binding them on both sides, rather than transporting cargo in dividing cells (Pandey et al., 2021; Walczak et al., 1997). During mitosis, KIF11 binds interpolar MTs and acts as a brake on other mitotic kinesins, ensuring proper spindle formation (Enos and Morris, 1990; Ferenz et al., 2010; Miki et al., 2001). Disruption in KIF11 function results in monopolar spindles and polyploidy (Kashina et al., 1996; Sharp et al., 1999). KIF11 binds both antiparallel MTs in the spindle and parallel astral MTs (Sharp et al., 1999). In vitro studies reveal that KIF11 slides antiparallel MTs but only crosslinks parallel MTs (Kapitein et al., 2005; Meißner et al., 2024). Although KIF11 is downregulated at the end of mitosis, it has also been found expressed in postmitotic neurons (Ferhat et al., 1998; Yount et al., 2015). Several studies have demonstrated roles for KIF11 in axon migration, branching, and dendritic arborization in maturing sympathetic and hippocampal neurons (Freixo et al., 2018; Kahn et al., 2015; Nadar et al., 2008). Despite the growing evidence of KIF11’s importance in developing neurons, the mechanism of its action in mature neurons and its potential contribution to MCLID remain unclear.

In a screen aimed at identifying MT motors involved in excitatory synaptic transmission in mouse primary hippocampal neurons, we unexpectedly found that Kinesin-5 (KIF11) acts as a repressor of excitatory synaptic transmission (Swarnkar et al., 2018). Specifically, knockdown of KIF11 in mature neurons led to an increase in the frequency of excitatory post-synaptic currents, enhanced dendritic arborization, and a higher number of dendritic spines. Additionally, we observed a decrease in KIF11 expression accompanied by elevated levels of synaptic vesicle proteins, such as Synaptophysin and Piccolo, within dendritic spines. Despite these findings, the mechanism by which KIF11 restrains dendritic growth and presynaptic vesicle release in mature neurons remains unclear. Given the structural and electrophysiological alterations resulting from KIF11 perturbations in primary hippocampal cultures, and the limited understanding of how KIF11 mutations lead to intellectual impairment in MCLID, we aimed to gain mechanistic insight into KIF11’s role in mature neurons.

Using live imaging of MT dynamics and neuronal cargoes in KIF11 knockdown neurons, we identified KIF11 as a crucial rheostat of MTs and intracellular transport in mature neurons. Pharmacological inhibition of KIF11 suggests that it stabilizes MTs by binding and crosslinking parallel, rather than antiparallel, MTs in mature neurons. Furthermore, the expression of two distinct MCLID-associated mutations linked to intellectual disabilities suggests that while these mutant forms of KIF11 retain their MT-binding capability, though with impaired sliding capacity, they negatively impact MT dynamics and dendritic complexity. Finally, the development of an optically controllable KIF11 revealed that temporal inhibition of KIF11 increases MT dynamics and dendritic growth, whereas temporal activation exerts force on MTs, diminishing both MT dynamics and dendritic growth. Collectively, these data establish KIF11 as a critical regulator of MT stability and dendritic arbor integrity in mature neurons. Additionally, they provide novel insights into the pathomechanism of MCLID, where mutant KIF11 proteins exert unrestrained forces on MTs, leading to their destabilization and a consequent reduction in dendritic arbor complexity.

## RESULTS

### KIF11 crosslinks, rather than slides MTs in mature neurons

To explore how KIF11 regulates dendritic arborization, we considered its known role in controlling cytoskeletal components, particularly MTs (MTs) in neurons, similar to KIF11-mediated stabilization of spindle MTs during cell division. In neurons, MTs are differentially organized: axons contain predominantly plus-end-out MTs, whereas dendrites exhibit a mixed polarity of MTs (Figure 1A).

**Figure 1.**
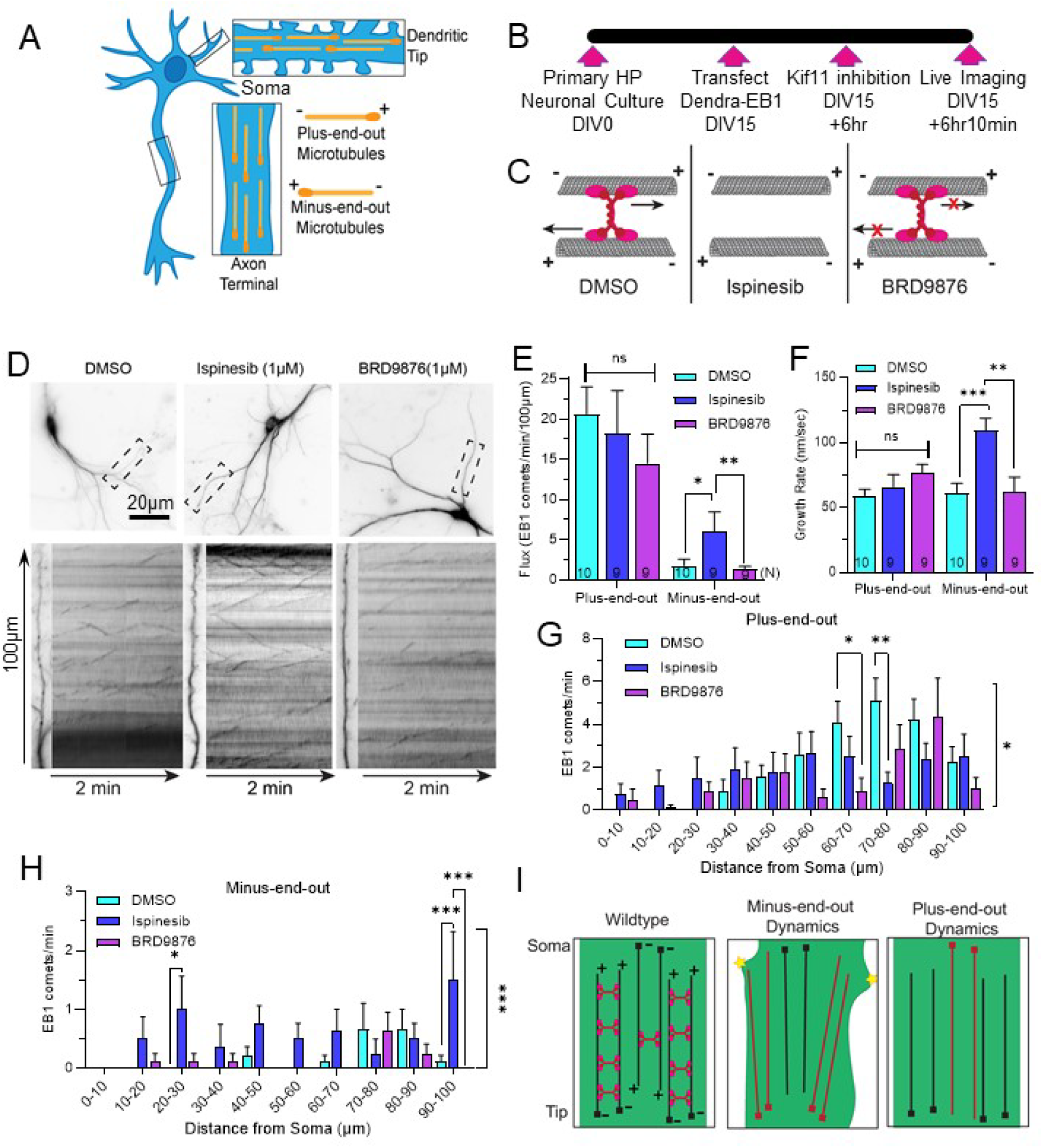
KIF11 crosslinks, rather than slide microtubules, in mature neurons. **A.** Schema microtubule orientation in neurons. Dendrites contain a mix of plus-end-out and minus-end-out microtubules. Axons primarily contain plus-end-out microtubules. **B.** Experimental timeline. **C**. Schema of how two different KIF11 inhibitors impact KIF11’s ability to bind microtubules. Ispinesib prevents KIF11 from binding microtubules. BRD9876 locks KIF11 on microtubules bound to the microtubule. **D.** Top: Primary hippocampal neurons expressing Dendra-EB1. The dashed line indicates the analyzed region. Bottom: Selected dendrites and their corresponding kymographs. **E**. EB1 comet flux from D for both anterograde (plus-end-out) and retrograde (minus-end-out) tracks. One-way ANOVA, Tukey’s multiple comparison test, *p-value<0.05, **p-value<0.01, ns=not significant. Error bars=SEM. **F**. EB1 comet growth rate from D for both anterograde (plus-end-out) and retrograde (minus-end-out) tracks. One-way ANOVA, Tukey’s multiple comparison test, *p-value<0.05, **p-value<0.01, ***p-value<0.001, Error bars=SEM. **G-H.** Distribution of anterograde (**G**) and retrograde (**H**) comets from D. Two-way ANOVA, Tukey’s multiple comparison test, * p-value <0.05, ** p-value<0.01, Error bars=SEM. **I.** Proposed model for KIF11 function in mature neurons. In Wildtype neurons, KIF11 stabilizes parallel microtubules. When this restriction is removed from the more exterior minus-end-out microtubules (red) their growth initiates new branch formation. When KIF11 is removed from the more interior plus-end-out microtubules this new growth (red) does not initiate new arborization.

We first confirmed the presence of KIF11 in the dendrites of mature neurons by detecting KIF11 mRNA and protein in hippocampal neurons (Supplementary Figure S1A,B). KIF11 has been shown to bind both parallel and antiparallel MTs, with its activity depending on the orientation of the MTs: KIF11 slides MTs when bound to antiparallel MTs and crosslinks them when bound to parallel MTs. To gain functional insights into whether KIF11 slides or crosslinks MTs in mature hippocampal neurons, we used two pharmacological inhibitors of KIF11 in primary hippocampal neurons expressing Dendra-EB1, a reporter for MT dynamics (Figure 1B). The inhibitors included Ispinesib, which prevents KIF11 from binding to MTs [24], and BRD9876, which locks KIF11 in a rigor state, bound to MTs [25] (Figure 1C). We hypothesized that if Ispinesib alone altered MT dynamics, KIF11 must be crosslinking MTs, whereas if BRD9876 had an effect, KIF11 was likely involved in sliding antiparallel MTs in mature dendrites.

We first evaluated MT flux and found that neither compound affected the plus-end-out MTs. However, Ispinesib treatment significantly increased the dynamics of minus-end-out MTs (Figure 1D-E, Supplementary Movie S1. Plus-end-out: DMSO = 20.667 ± 3.362 #/100µm/min, Ispinesib = 18.250 ± 5.324 #/100µm/min, BRD9876 = 14.500 ± 3.650 #/100µm/min; Minus-end-out: DMSO = 1.778 ± 0.778 #/100µm/min, Ispinesib = 6.000 ± 2.435 #/100µm/min, BRD9876 = 1.250 ± 0.412 #/100µm/min. N=10-9 neurons; Two-Way ANOVA followed by Tukey’s; *p<0.05, **p<0.01). Next, we assessed the MT growth rate and observed that while neither compound affected plus-end-out MTs, Ispinesib treatment led to a significant increase in the growth rate of minus-end-out MTs (Figure 1F. Plus-end-out: DMSO = 50.305 ± 4.978 nm/sec, Ispinesib = 66.084 ± 9.392 nm/sec, BRD9876 = 77.124 ± 6.082 nm/sec; Minus-end-out: DMSO = 61.192 ± 7.432 nm/sec, Ispinesib = 109.544 ± 9.269 nm/sec, BRD9876 = 61.916 ± 11.681 nm/sec. N=10-9 neurons; Two-Way ANOVA followed by Tukey’s; **p<0.01, ***p<0.001).

We further examined how MT dynamics were affected along the dendrites. Ispinesib treatment resulted in a decrease in plus-end-out EB1 comets in distal dendrites (60-70µm from the soma) and an increase in minus-end-out EB1 comets in proximal dendrites (20-30µm and 90-100µm from the soma). In contrast, BRD9876 treatment only decreased plus-end-out EB1 comets at 60-70µm from the soma (Figure 1G-H. Two-Way ANOVA followed by Tukey’s; *p<0.05, **p<0.01, ***p<0.001). These data suggest that KIF11 limits the presence of minus-end-out MTs by crosslinking and limiting growth of parallel MT bundles in mature neurons (Figure 1I).

### KIF11 mediates EB3 dynamics in mature neurons

To confirm the role of KIF11 in regulating MT (MT) dynamics within mature neurons, we investigated MT behavior in KIF11 knockdown neurons. We transfected primary hippocampal neurons with a far-red-tagged end-binding protein 3 (miRFP703-EB3) to track growing MTs, alongside either a negative control construct (NC-GFP) or a previously validated shRNA construct targeting KIF11 (shKIF11) [23]. Imaging was conducted approximately 24 hours post-transfection (Figure 2A). Using this approach, we visualized EB3 comets, representing both plus- and minus-end-out MTs, in the axons and dendrites of NC-GFP and shKIF11 neurons (Figure 2B-E, Supplementary Movie S2).

**Figure 2.**
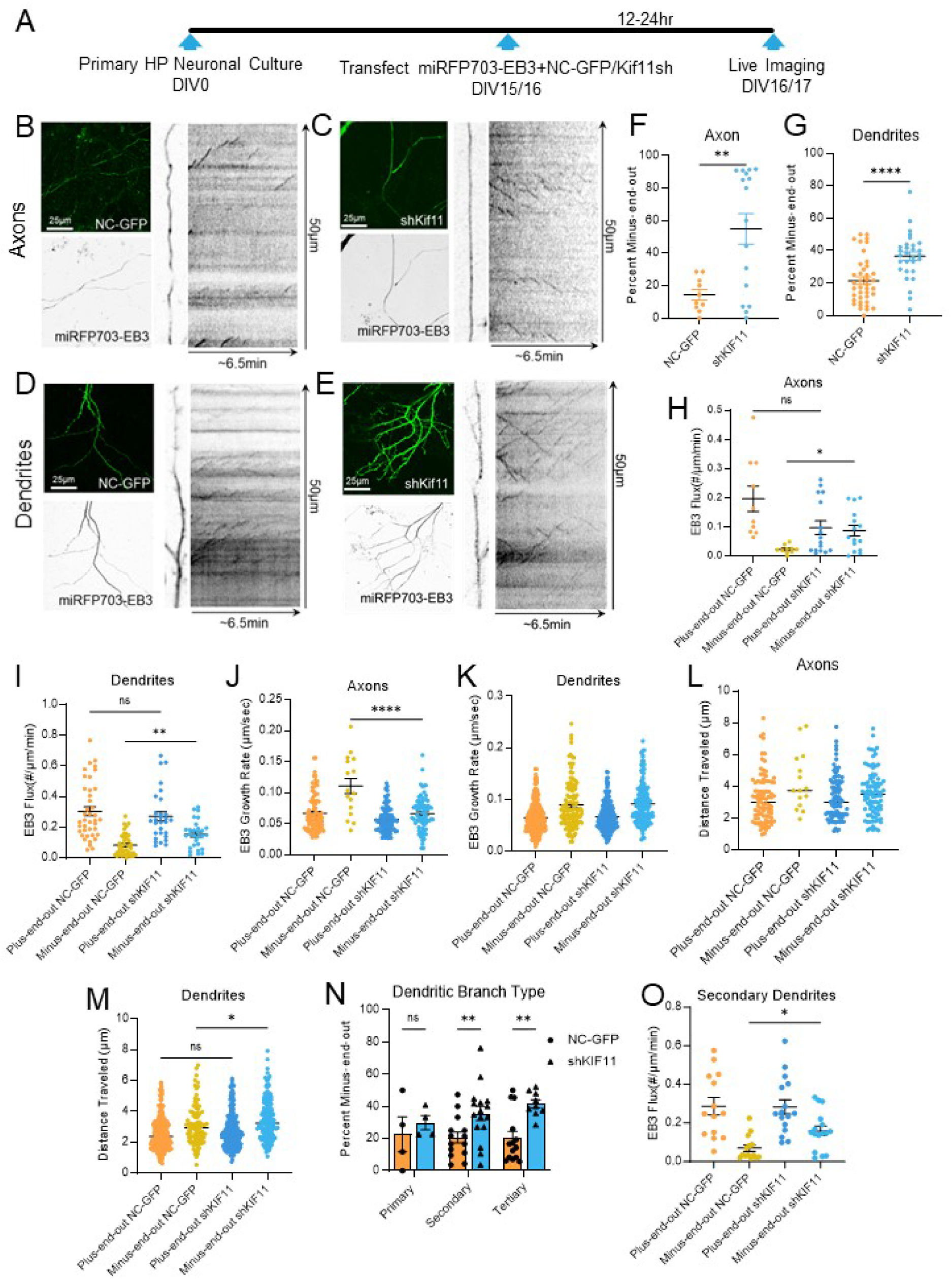
KIF11 regulates EB3 dynamics in mature neurons. **A**. Experimental timeline. **B.** Left: Maximum projection intensity images of an axon from cells co-expressing NC-GFP (scrambled negative control) and miRFP703-EB3. Right: Selected axon and kymograph of miRFP703-EB3. **C**. Left: Maximum projection intensity images of axons from cells co-expressing shKif11 (shRNA targeting KIF11) and miRFP703-EB3. Right: Selected axon and kymograph of miRFP703-EB3 **D.** Left: Maximum projection intensity images of dendrites from cells co-expressing NC-GFP (scrambled negative control) and miRFP703-EB3. Right: Selected dendrite and kymograph of miRFP703-EB3. **E**. Left: Maximum projection intensity images of dendrites from cells co-expressing shKif11 (shRNA targeting KIF11) and miRFP703-EB3. Right: Selected dendrite and kymograph of miRFP703-EB3. **F, G.** The percentage of minus-end-out microtubules in axons (**F**) and dendrites (**G**) in NC-GFP or shKIF11 neurons. Unpaired t-test, **p-value<0.01, ****p-value<0.0001. Error bars=SEM. **H, I.** EB3-comet flux in axons (**H**) and dendrites (**I**) in NC-GFP or shKIF11 neurons. One-way ANOVA, Tukey’s multiple comparison test, **** p-value<0.0001, Error bars=SEM. **J, K.** EB3-comet growth rate in axons (**J**) and dendrites (**K**) in NC-GFP or shKIF11 neurons. One-way ANOVA, Tukey’s multiple comparison test, * p-value<0.05, p-value<0.01, ns=not significant. Error bars=SEM. **L, M.** EB3-comet distance traveled (MT growth) in axons (**L**) and dendrites (**M**) in NC-GFP or shKIF11 neurons. One-way ANOVA, Tukey’s multiple comparison test, * p-value<0.05, ns=not significant. Error bars=SEM. **N.** The percentage of minus-end-out microtubules in primary, secondary, and tertiary dendrites. Mixed-effects model (REML) followed by Tukey’s post hoc test. **p<0.01, Error bars= SEM. **O.** EB3-comet flux for plus-end-out and minus-end-out EB3 comets in secondary dendrites. One-way ANOVA, Tukey’s multiple comparison test, *p-value <0.05. Error bars=SEM.

Our results revealed a significant increase in the percentage of minus-end-out MTs in shKIF11 axons compared to control neurons (Figure 2F. NC-GFP = 14.30±3.24%; shKIF11 = 54.85±9.36%; N = 10-16 axons, **p<0.01). A similar increase was observed in shKIF11 dendrites (Figure 2G. NC-GFP = 21.48±2.27%; shKIF11 = 36.62±2.67%; N = 40-29 dendrites, **p<0.01, Brown-Forsythe ANOVA). Additionally, KIF11 knockdown led to an increase in minus-end-out MT flux in both axons and dendrites (Figure 2H, I. H: plus-end-out NC-GFP = 0.197±0.043 #/µm/min; minus-end-out NC-GFP = 0.023±0.004 #/µm/min; plus-end-out shKIF11 = 0.097±0.024 #/µm/min; minus-end-out shKIF11 = 0.087±0.018 #/µm/min, N = 10-16 axons, *p<0.05. I: plus-end-out NC-GFP = 0.305±0.029 #/µm/min; minus-end-out NC-GFP = 0.083±0.011 #/µm/min; plus-end-out shKIF11 = 0.272±0.030 #/µm/min; minus-end-out shKIF11 = 0.154±0.017 #/µm/min, **p<0.01, N = 40-29 dendrites, Brown-Forsythe ANOVA).

We further assessed MT growth rates, discovering a decrease in the minus-end-out EB3 growth rate in axons, while no significant change was observed in dendrites (Figure 2J, K. J: plus-end-out NC-GFP = 0.067±0.003 µm/sec; minus-end-out NC-GFP = 0.111±0.012 µm/sec; plus-end-out shKIF11 = 0.057±0.002 µm/sec; minus-end-out shKIF11 = 0.066±0.003 µm/sec, N = 10-16 axons, ****p<0.0001. K: plus-end-out NC-GFP = 0.065±0.001 µm/sec; minus-end-out NC-GFP = 0.090±0.004 µm/sec; plus-end-out shKIF11 = 0.066±0.002 µm/sec; minus-end-out shKIF11 = 0.092±0.003 µm/sec, N = 40-29 dendrites, Brown-Forsythe ANOVA).

When examining the distance traveled by EB3, which reflects MT growth, no significant change was observed in axons (Figure 2L. plus-end-out NC-GFP = 3.250±0.162 µm; minus-end-out NC-GFP = 4.285±0.467 µm; plus-end-out shKIF11 = 3.247±0.145 µm; minus-end-out shKIF11 = 3.524±0.158 µm, N = 10-16 axons, Brown-Forsythe ANOVA). However, a significant increase in the growth of minus-end-out MTs was detected in dendrites (Figure 2M. plus-end-out NC-GFP = 2.551±0.054 µm; minus-end-out NC-GFP = 3.077±0.071 µm; plus-end-out shKIF11 = 2.739±0.071 µm; minus-end-out shKIF11 = 3.502±0.106 µm, N = 40-29 dendrites, Brown-Forsythe ANOVA, *p<0.05). Collectively, these data suggest that KIF11 normally constrains minus-end-out MT dynamics in both axons and dendrites, thereby limiting arborization.

To determine whether these alterations in MT dynamics were specific to certain types of dendrites or distributed throughout the entire arbor, we conducted a segmented analysis of EB3 dynamics using the dataset from Figure 2. We observed an increase in the percentage of dynamic minus-end-out MTs in secondary and tertiary dendrites, but not in primary dendrites (Figure 2N. Primary NC-GFP = 22.584±10.857% N = 4, Primary shKIF11 = 29.888±4.454% N = 4; Secondary NC-GFP = 20.650±3.430% N = 14, Secondary shKIF11 = 35.513±4.408% N = 16; Tertiary NC-GFP = 20.228±4.185% N = 14, Tertiary shKIF11 = 41.592±2.534% N = 9 dendrites; Mixed-effects model (REML), **p<0.01). Further analysis revealed a significant increase in minus-end-out EB3 flux specifically in secondary shKIF11 dendrites (Supplementary Figure S2 A-H; Figure 2O. plus-end-out NC-GFP = 0.287±0.045 #/µm/min; minus-end-out NC-GFP = 0.070±0.017 #/µm/min; plus-end-out shKIF11 = 0.283±0.037 #/µm/min; minus-end-out shKIF11 = 0.159±0.025 #/µm/min, NC-GFP N = 14, shKIF11 N = 16 dendrites, *p<0.05, Brown-Forsythe ANOVA). This indicates that KIF11 specifically restricts the number of growing minus-end-out MTs within dendrites, particularly in secondary and tertiary branches, thereby limiting the formation of new branches.

### KIF11 regulates the amount of mobile and stationary synaptophysin in neurons

Our previous data demonstrate that functional depletion of KIF11 leads to significant destabilization of dendritic MTs (MTs). We next investigated whether this instability would affect the transport of cargos that rely on these MTs for movement. Previous work from our lab revealed that KIF11 depletion resulted in the accumulation of synaptophysin without affecting PSD95 levels, so we focused our analysis on these two MT motor cargos (Swarnkar et al., 2018).

To examine synaptophysin dynamics, we co-transfected mature hippocampal neurons with mRuby-Synaptophysin and either NC-GFP or KIF11sh constructs (Figure 3A, Supplementary Movie S3). KIF11 knockdown led to a decrease in the motility of mRuby-synaptophysin compared to the NC-GFP control (Figure 3B-D. NC-GFP= 46.32±2.31%, N=27 processes; shKIF11=56.19±2.60%, N=38 processes; Unpaired Student’s T-test, **p<0.01). However, the distance transported, velocity, and flux of mRuby-Synaptophysin were not affected by KIF11 knockdown (Figure 3B-C, E-G). These results suggest that impaired KIF11 function causes synaptophysin accumulation in neurons by reducing the proportion of mobile synaptophysin, rather than altering the velocity or flux of the moving molecules.

**Figure 3.**
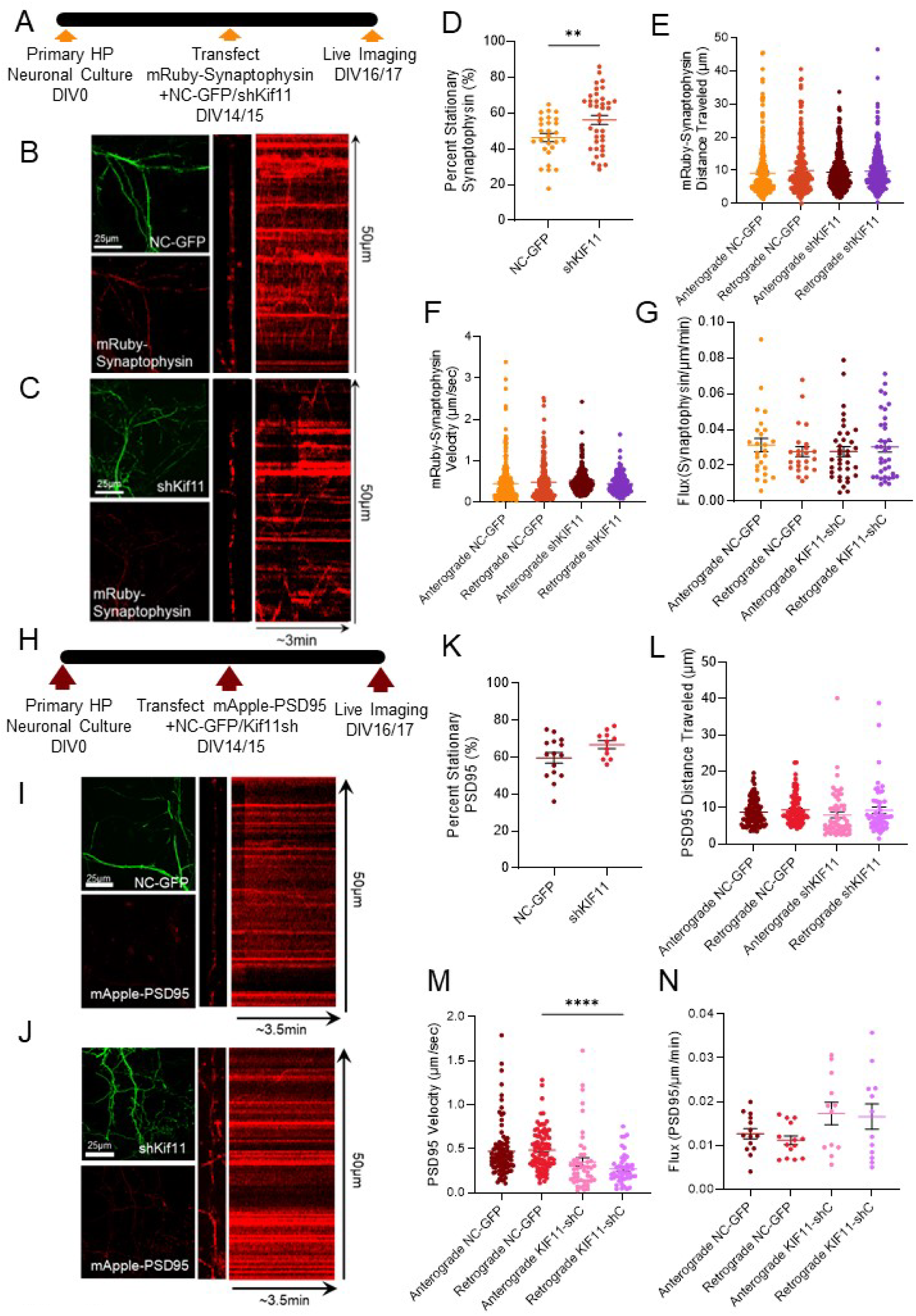
KIF11 regulates the amount of mobile and stationary synaptophysin in neurons. **A.** Experimental timeline. **B.** Left: Maximum projection intensity images of dendrites from cells co-expressing NC-GFP (scrambled negative control) and mRuby -Synaptophysin Right: Selected dendrite and kymograph of mRuby-Synaptophysin. **C**. Left: Maximum projection intensity images of dendrites from cells co-expressing shKif11 (shRNA targeting KIF11) and mRuby-Synaptophysin. Right: Selected dendrite and kymograph of mRuby-Synaptophysin. **D.** Percent of stationary mRuby-Synaptophysin in NC-GFP and KIF11-KD primary hippocampal neurons. Unpaired t-test, **p-value<0.01, Error bars=SEM. **E, F, G.** The distance traveled by mRuby-Synaptophysin (**E**), the velocity of mRuby-Synaptophysin (**F**), and the Flux of mRuby-Synaptophysin (**G**) in NC-GFP and KIF11-KD primary hippocampal neurons. One-way ANOVA, Tukey’s multiple comparison test. Error bars=SEM. **H.** Experimental timeline. **I.** Left: Maximum projection intensity images of dendrites from cells co-expressing NC-GFP (scrambled negative control) and mApple-PSD95 Right: Selected dendrite and kymograph of mApple-PSD95. **J**. Left: Maximum projection intensity images of dendrites from cells co-expressing shKif11 (shRNA targeting KIF11) and mApple-PSD95. Right: Selected dendrite and kymograph of mApple-PSD95 **K.** Percent of stationary mApple-PSD95 in NC-GFP and KIF11-KD primary hippocampal neurons. Unpaired t-test. Error bars=SEM. **L, M, N.** The distance traveled by mApple-PSD95 (**L**), the velocity of mApple-PSD95 (**M**), and the Flux of mApple-PSD95 (**N**) in NC-GFP and KIF11-KD primary hippocampal neurons. One way ANOVA. Tukey’s multiple comparison test. ****p-value<0.001. Error bars=SEM.

Next, we assessed PSD95 dynamics by co-transfecting mature hippocampal neurons with mApple-PSD95 and either NC-GFP or KIF11sh constructs (Figure 3H, Supplementary Movie S4). We observed no change in the percentage of stationary mApple-PSD95 trafficking in KIF11 knockdown neurons (Figure 3I-K). Additionally, KIF11 depletion did not affect the distance traveled or the flux of mApple-PSD95 (Figure 3I-J, L-M). Although KIF11 depletion reduced the retrograde velocity of mApple-PSD95, this did not lead to PSD95 accumulation (Figure 3N). Consistent with our previous findings, KIF11 depletion does not alter PSD95 abundance in dendrites.

### Transient KIF11 inhibition disrupts the transport of both mitochondria and lysosomal vesicles

We next investigated whether functional depletion of KIF11 affects intracellular trafficking beyond synaptic proteins. To address this, we examined the transport of key organelles—mitochondria and lysosomal vesicles (LVs)—in mature neurons. Mitochondria and LVs are transported by various MT motors on both stabilized and dynamic MTs, making them ideal targets for assessing broader transport alterations.

To gain functional insights, we pharmacologically inhibited KIF11 using Ispinesib and BRD9876 (Supplementary Figure S3A). We labeled mitochondria and LVs in DIV16 hippocampal neurons with MitoTracker Green and LysoTracker Red, respectively, to ensure complete labeling (Supplementary Figure S3B, Supplementary Movie S5).

Our analysis of mitochondrial transport revealed that Ispinesib treatment decreased anterograde flux and increased retrograde flux, whereas BRD9876 did not produce significant changes compared to the DMSO control (Supplementary Figure S3C-D). Given that BRD9876 did not significantly impact our previous analyses, we proceeded to evaluate LVs using only DMSO and Ispinesib. Ispinesib treatment led to a trend of increased anterograde flux of LVs and significantly increased retrograde flux (Supplementary Figure S3E-F, Supplementary Movie S6). Together, these findings indicate that KIF11 inhibition significantly disrupts the transport of both mitochondria and LVs in mature hippocampal neurons.

### Intellectual disability causing mutations in KIF11 reduce the complexity of dendritic arborization and alter MT dynamics

To explore how KIF11 gain-of-function affects dendritic arborization and MT (MT) dynamics, and to assess the impact of KIF11 mutations associated with severe intellectual disabilities and minor microcephaly, we focused on two specific mutations: Y82F (Y81F in Mus musculus) and H768Qfs*7 (represented as KIF11^ΔCterm^) (Figure 4A) (Schlögel et al., 2015).

**Figure 4.**
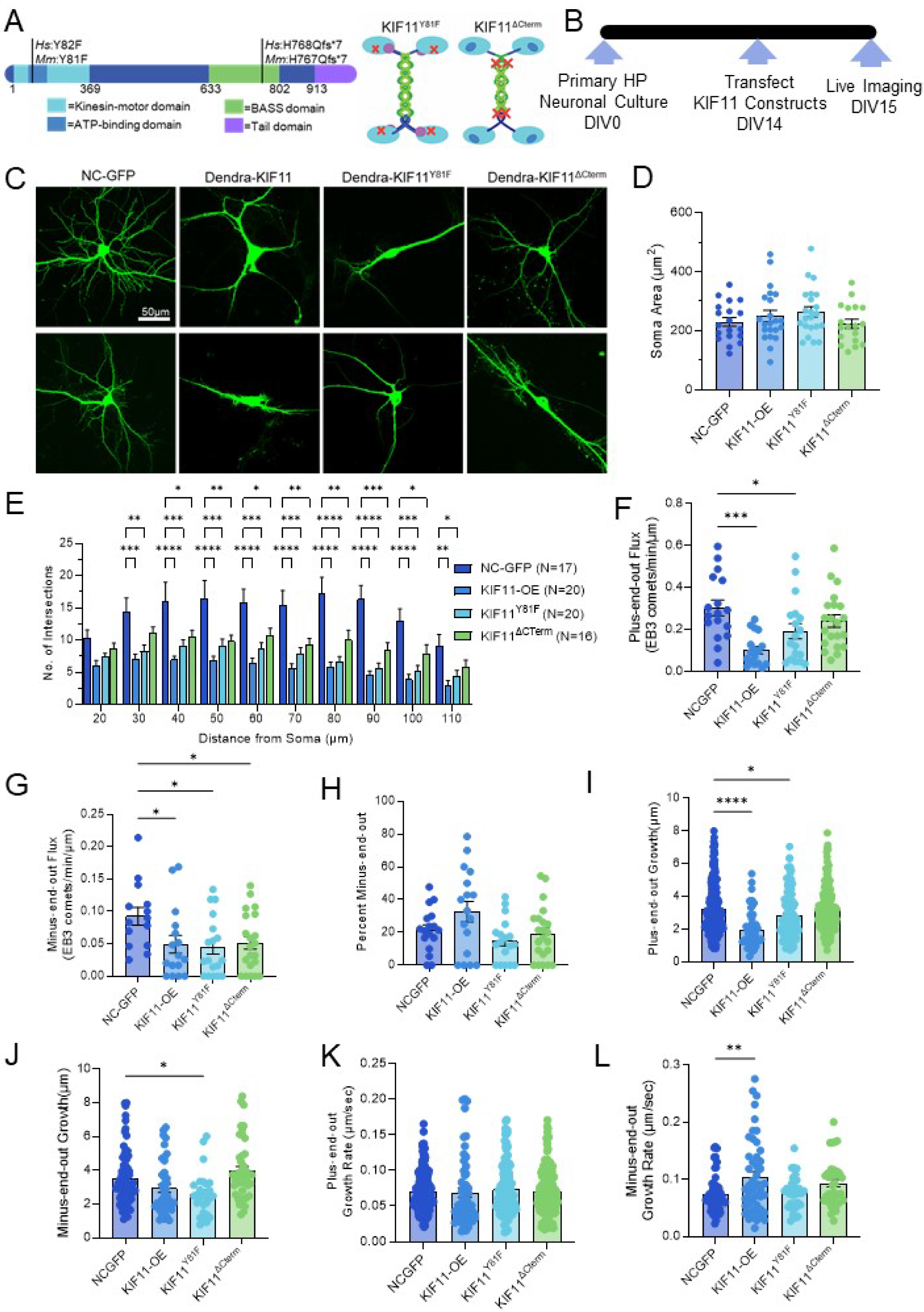
KIF11 patient mutations impact dendritic arborization and microtubule dynamics. **A.** Schema of KIF11 and locations of selected Microcephaly with or without chorioretinopathy, lymphedema, or intellectual disabilities syndrome (MCLID) patient mutations (*Hs: Homo sapiens*) and the corresponding mouse homolog (Mm: *Mus Musculus*) (adapted from Schlögel et al., 2015). **B**. Experimental timeline. **C.** Confocal projection images of primary hippocampal mouse neurons transfected with control or MCLID patient mutation constructs with soma in the center to depict the dendritic arbor. Scale Bar=25µm. **D.** Soma size quantification of (C). N=20,22,22,19 neurons for NCGFP, KIF11-OE, KIF11^Y81F^, and KIF11^ΔCterm^, respectively. One-way ANOVA followed by Tukey’s post hoc test. Error bars= SEM. **E**. Quantification of dendritic morphology changes using Sholl analysis of intersections per 10-µm step size. N=17,20,20,16 neurons for NCGFP, KIF11-OE, KIF11^Y81F,^ and KIF11^ΔCterm^, respectively. Two-way ANOVA followed by Tukey’s post hoc test. Error bars= SEM. *p<0.05, **p<0.01. **F.** Plus-end-out EB3-comet flux in KIF11 and KIF11 mutant dendrites in comparison to NC-GFP. N=17, 17,18,21 dendrites for NCGFP, KIF11-OE, KIF11^Y81F^, and KIF11^ΔCterm^, respectively. One-way ANOVA followed by Dunnett’s test, *p<0.05, ***p<0.001. Error bars=SEM. **G.** Minus-end-out EB3-comets flux in KIF11 and KIF11 mutant dendrites in comparison to NC-GFP. N=14,16,18,21 dendrites for NCGFP, KIF11-OE, KIF11^Y81F^, and KIF11^ΔCterm^, respectively. One-way ANOVA followed by Dunnett’s test. Error bars=SEM. **H.** Percentage of Minus-end-out EB3-comets in KIF11 and KIF11 mutant dendrites in comparison to NC-GFP. N=17,17,18,22 dendrites for NCGFP, KIF11-OE, KIF11^Y81F^, and KIF11^ΔCterm^, respectively. One-way ANOVA followed by Dunnett’s test. Error bars=SEM. **I.** Amount of plus-end-out microtubule growth based on distance traveled by EB3-comets in KIF11 and KIF11 mutant dendrites in comparison to NC-GFP. N=17(248),17(59),18(114),22(184) dendrites (# of comets) for NCGFP, KIF11-OE, KIF11^Y81F^, and KIF11^ΔCterm^, respectively. One-way ANOVA followed by Dunnett’s test, *p<0.05, ***p<0.001. Error bars=SEM. **J.** Amount of minus-end-out microtubule growth based on distance traveled by EB3-comets in KIF11 and KIF11 mutant dendrites in comparison to NC-GFP. N=17(70),17(42),18(27),22(45) dendrites (# of comets) for NCGFP, KIF11-OE, KIF11^Y81F^, and KIF11^ΔCterm^, respectively. One-way ANOVA followed by Dunnett’s test, *p<0.05. Error bars=SEM. **K.** Plus-end-out microtubule growth rate based on EB3-comet velocities in KIF11 and KIF11 mutant dendrites in comparison to NC-GFP. N=17(254),17(56),18(130),22(189) dendrites (# of comets) for NCGFP, KIF11-OE, KIF11^Y81F^, and KIF11^ΔCterm^, respectively. One-way ANOVA followed by Dunnett’s test. Error bars=SEM. **L.** Minus-end-out microtubule growth rate based on EB3-comet velocities in KIF11 and KIF11 mutant dendrites in comparison to NC-GFP. N=17(59),17(49),18(32),22(41) dendrites (# of comets) for NCGFP, KIF11-OE, KIF11^Y81F^, and KIF11^ΔCterm^, respectively. One-way ANOVA followed by Dunnett’s test, **p<0.01. Error bars=SEM.

We began by examining dendritic arborization. Primary hippocampal neurons were transfected with either a control construct (NC-GFP), KIF11 overexpression construct (CAG-Dendra-KIF11), or constructs carrying the KIF11^Y81F^ or KIF11^ΔCterm^ murine versions of patient mutations. Neurons were imaged approximately 24 hours later to assess dendritic morphology (Figure 4B-C). Although these mutations are linked with microcephaly, no significant difference was observed in soma area among the conditions (Figure 4D). Sholl analysis demonstrated that KIF11 overexpression, as well as the KIF11^Y81F^ or KIF11^ΔCterm^ mutations, led to a marked reduction in dendritic complexity, with the most severe effects observed in KIF11-OE and KIF11^Y81F^ (Figure 4E). These findings further confirm the essential role of KIF11 in regulating dendritic arborization.

Given that KIF11 depletion increases MT dynamics, we hypothesized that KIF11 overexpression might decrease MT dynamics. To test this, we co-transfected neurons with miRFP703-EB3 and either NC-GFP, KIF11-OE, or the mutant constructs and imaged them ∼24 hours later (Supplementary Figure S4A-E, Supplementary Movie S7). We observed a significant decrease in the flux of plus-end-out MTs in KIF11-OE and KIF11^Y81F^ neurons, with a slight decrease noted in KIF11^ΔCterm^ neurons (Figure 4F). Additionally, a trend towards reduced flux of minus-end-out MTs was seen in KIF11-OE and mutant neurons (Figure 4G). Although there were no statistically significant differences in the percentage of minus-end-out MTs (Figure 4H), a reduction in the distance of MT growth in both plus- and minus-end-out directions was observed for KIF11-OE and KIF11^Y81F^ neurons (Figure 4I,J). No differences were detected in the growth rate of plus-end-out MTs (Figure 4K). However, KIF11-OE neurons exhibited an increased growth rate for minus-end-out MTs, suggesting that the few remaining growing microtubules are destablilized (Figure 4I). Overall, KIF11 overexpression and the KIF11^Y81F^ mutation significantly constrain MT growth dynamics.

During MT dynamics imaging in KIF11-OE and KIF11 mutant neurons, we noted numerous bent, twisted, and looped MTs (Supplementary Figure 4F-H). Previous in vitro studies suggested that KIF11 might act as an MT polymerizer, producing curved and looped MTs during polymerization (Chen and Hancock, 2015). However, we observed that these loops often lacked EB3 comets and were not initiated by EB3 comets over more than 8 hours of imaging. These observations imply that the loops are not the result of new MT growth. Instead, we captured the process of MTs bending, twisting, and forming loops on themselves (Supplementary Figure 4I, Supplementary Movie S8). This twisting likely reflects a combination of torque and tension applied to the MTs (Krieg et al., 2017). Additionally, we observed dead and dying neurons in KIF11-OE and KIF11^Y81F^ cultures with bent and fragmented dendrites, indicating that these MTs eventually succumb to the applied tension and break (Supplementary Figure 3J-K). These findings align with the concept that KIF11 exerts rotational force when crosslinking antiparallel/parallel MTs, as seen in vitro with full-length KIF11 in a 3D MT sliding assay (Meißner et al., 2024). In summary, the observation of twisted and looped MTs in KIF11 overexpression conditions supports the role of KIF11 as an MT crosslinker in neurons.

### MCLID mutations differentially impact KIF11 velocity

While we observed that both KIF11^Y81F^ and KIF11^ΔCterm^ reduced dendritic complexity and minus-end-out MT flux, similar to WT KIF11, there were several notable differences (Figure 4C-E,G). Neither mutant increased minus-end-out MT growth rate (Figure 4L), KIF11^Y81F^ more significantly decreased minus-end growth (Figure 4J), and KIF11^ΔCterm^ allowed visualization of spines (Figure 4C, Supplementary Figure 4H) suggestive of diffusion. These observations suggest that while these mutants still retain microtubule binding capabilities, KIF11 motility is likely impaired.

The residue Y82 is positioned between the ATP binding pocket and the tail binding domain (Figure 5A). Considering the importance of the coordination of tail association in the mechanochemical cycle of the motor domain, the conservative mutation Y82F may be enough to disrupt it (Bodrug et al., 2020). Additionally, H768 residue, located within the bilateral assembly (BASS) domain has been identified to participate in the packing of antiparallel helices (Scholey et al., 2014). Thus, the KIF11^ΔCterm^ mutant is both mutated within the BASS domain and truncated before the tail domain, both the formation of the tetramer and the coordination of the mechanochemical cycling could be affected (Figure 5B). We performed a microtubule sliding assay using synthesized full-length WT human KIF11, KIF11^Y82F^, and KIF11^ΔCterm^ to investigate the impact these mutations have on KIF11’s MT binding more directly (Figure 5C). Interestingly, KIF11^Y82F^ was found to significantly decrease KIF11 velocity while KIF11^ΔCterm^ increased it and the distribution of velocities (Figure 5D). These findings show that these mutants still retain MT binding capabilities but display a loss-of-function and gain-of-function respectively.

**Figure 5.**
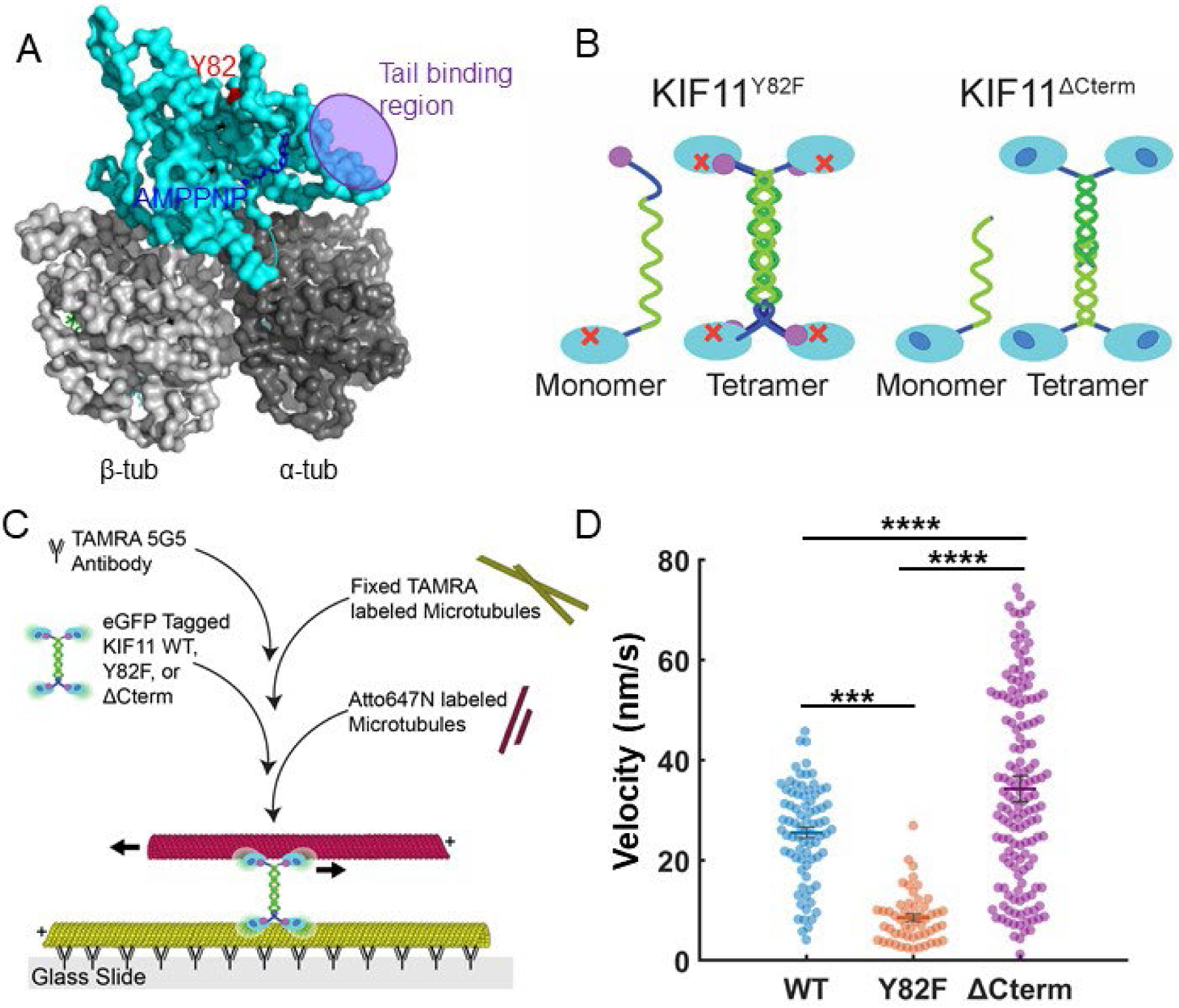
Intellectual disability-causing mutations in KIF11 impact microtubule sliding capacity. **A.** Surface plot diagram based on Protein Data Bank (PDB)-ID: 6VPO KIF11 motor (teal), AMPPNP (dark blue), and MT (grey). The Y82 residue is highlighted in red and the tail binding region is highlighted in purple (Bodrug et al., 2020). **B.** Models of human KIF11 proteins with intellectual disability-causing mutation. **C.** Schema of 2D microtubule sliding assay. **D.** Effect of KIF11^Y82F^ and KIF11^ΔCterm^ mutations on microtubule sliding velocity. N=53, 78, and 144 microtubules each for human KIF11, KIF11^Y82F^, and KIF11^ΔCterm^, respectively. Unpaired students t-test, ***p-value<0.001, ****p-value<0.0001, Error bars=SEM

### Local inhibition of KIF11 stimulates new dendritic growth

Our data reveal that KIF11 depletion increases MT (MT) dynamics and dendritic complexity, whereas KIF11 overexpression reduces both MT dynamics and dendritic complexity. These effects suggest that KIF11 plays a significant role in regulating MTs and dendritic arborization on a global scale. However, it remains unclear whether these changes result from KIF11’s direct or indirect impact on MTs and dendritic structure.

To better elucidate KIF11’s role in mature neurons, we sought to locally inhibit KIF11 and assess its effects on MT dynamics and dendritic branching. We employed the pdDronpaV system for this purpose. The pdDronpaV system allows for reversible dimerization of pdDronpaV molecules upon exposure to violet (405 nm) light and dissociation upon exposure to cyan (488 nm) light (Zhou et al., 2017). By tagging KIF11 at the N-terminus with pdDronpaV, we hypothesized that the monomeric form of pdDronpaV would not affect KIF11’s function, similar to Dendra-KIF11 (Figure 5A). However, dimerization of pdDronpaV would distort the motor and tail domains of KIF11, impairing its ability to bind MTs effectively (Figure 5A).

We first tested the dimerization capability of pdDronpaV within the KIF11 tetramer by transfecting HEK293T cells with either pdDronpaV-CDK5 or pdDronpaV-KIF11, along with miRFP703-EB3. We observed that exposure to cyan light reduced the fluorescence of pdDronpaV-CDK5, while violet light increased the fluorescence intensity of pdDronpaV-KIF11, indicating successful dimerization (Figure 5B-D, Supplementary Movie S9). Importantly, neither cyan nor violet light affected miRFP703-EB3 fluorescence, suggesting that changes in pdDronpaV fluorescence were due to dimerization rather than alterations in EB3 expression or fluorescence.

Next, we investigated the effects of temporally inhibiting KIF11 in primary hippocampal neurons by co-expressing pdDronpaV-KIF11 and miRFP703-EB3. Inhibition of pdDronpaV-KIF11 with violet light for 5 minutes resulted in a significant increase in the flux of growing plus-end-out (anterograde) and minus-end-out (retrograde) MTs (Figure 5D-F, Supplementary Movie S10). This increased MT growth was accompanied by enhanced dendritic growth (Figure 5F,H). Collectively, these findings demonstrate that KIF11 functions as a repressor of MT dynamics and dendritic growth in mature neurons.

### Local activation of KIF11 results in twisted MTs and dendritic branches

Our observations revealed that prolonged overexpression (>24 hours) of KIF11 resulted in twisted and compressed MTs (MTs), prompting us to investigate how KIF11 might exert torque on these structures. Specifically, we sought to understand whether this might provide insight into the orientation of KIF11 binding to MTs [30]. Pharmacological inhibition studies (Figure 1) indicated that KIF11 binds to parallel MT bundles. However, the large loops and contraction of dendritic arbors observed with KIF11 overexpression (Supplementary Figure S4F-I) suggest that KIF11 may also interact with anti-parallel MTs, causing rotation, whereas smaller kinks and loops are more characteristic of parallel binding (Figure 6A).

**Figure 6.**
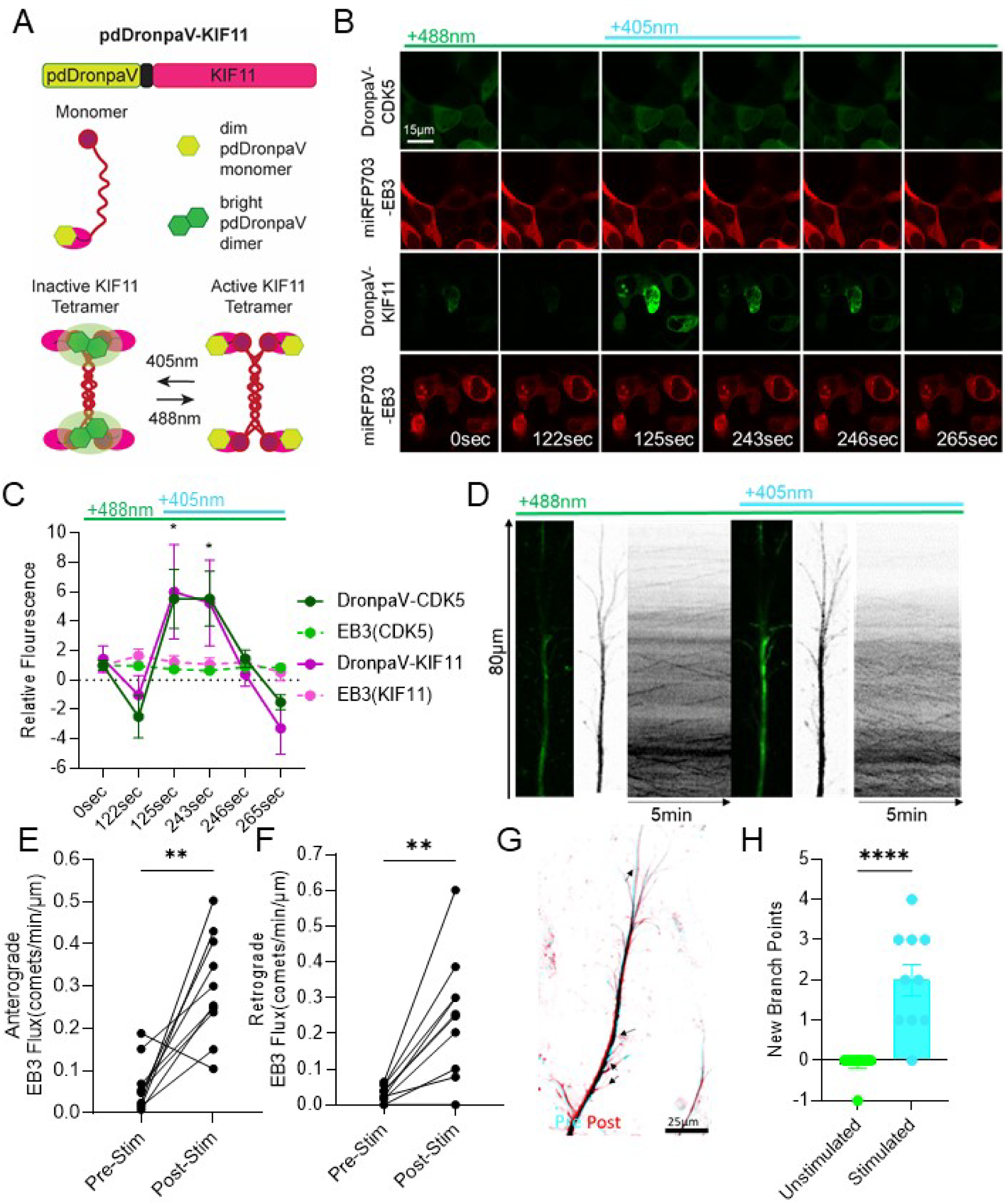
Local inhibition of KIF11 stimulates new dendritic growth. **A.** Schema of pdDronpaV-KIF11. 405nm stimulation leads to enhanced pdDronpaV fluorescence and the inactivation of KIF11. 488nm stimulation depletes pdDronpaV fluorescence and activates KIF11. **B.** Single frames of movies of HEK293T cells expressing the control (pdDronpaV-CDK5) or pdDronpaV-KIF11 and miRFP703-EB3.The duration of stimulation or inhibition is displayed at the top. **C.** Relative fluorescence intensity values from B calculated from Corrected Total Cell Fluorescence. N=10 cells all conditions. Two-way ANOVA, Tukey’s multiple comparison test, *p-value<0.05, Error bars=SEM. **D.** Example Dendrite from DIV17 primary hippocampal neurons expressing pdDronpaV-KIF11(left) and miRFP703(right) and the corresponding kymographs. Light wavelength stimulation shown at the top. **E,F.** Anterograde(E) and Retrograde (F) EB3 flux from pdDronpaV-KIF11 neurons pre- and post-stimulation with violet light. N=10 cells all conditions. Paired students t-test, **p-value<0.01, Error bars=SEM. **G.** Example Dendrite showing new branch points(arrowheads point to growth longer than 5µm) before(blue) and 10 minutes after stimulation (red) with violet light. **H**. Quantification of G. N=10 cells all conditions. Un-paired students t-test, ****p-value<0.0001, Error bars=SEM

**Figure 7.**
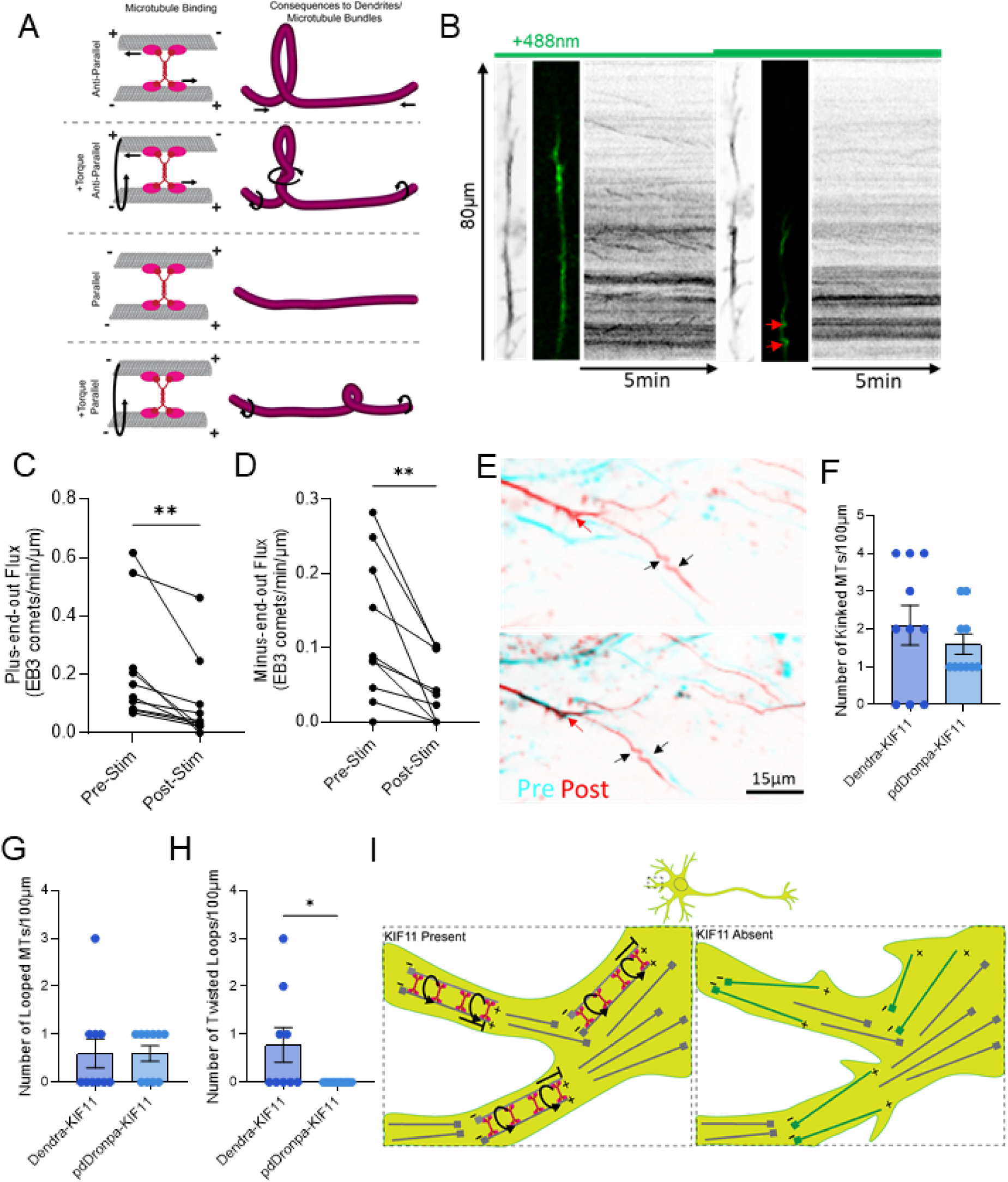
KIF11 enacts torque on parallel microtubule bundles in mature neurons. **A.** Schema of how rotational and non-rotational forces KIF11 enacts on microtubules, and the orientation in which it binds may result in twisted or looped dendrites. **B.** Activation of pdDronpa-KIF11 with cyan light results in kinked microtubules and decreased EB3 dynamics. A representative dendrite from DIV17 primary hippocampal neurons expressing pdDronpaV-KIF11(left) and miRFP703(right) and the corresponding kymographs. Light wavelength stimulation is shown at the top. Red arrows point to kinked microtubules. **C,D.** Anterograde(C) and Retrograde (D) EB3 flux from pdDronpaV-KIF11 neurons pre- and post-stimulation with cyan light. N=10 neurons. Paired students t-test, **p-value<0.01, Error bars=SEM. **E.** Example Dendrite showing kinked (black arrows) and looped (red arrow) microtubules before(blue) and 10 minutes after stimulation (red) with cyan light. **E,F,G**. Temporal activation of KIF11 (pdDronpa) results in looped and kinked MTs, suggesting parallel binding. Constitutive activation of KIF11 (Dendra-KIF11) results in looped, kinked, and twisted looped MTs (see Supp. Fig4I) suggesting parallel and anti-parallel binding to microtubules. Quantification of E. N=10 cells. Unpaired students t-tests, *p-value<0.05, Error bars=SEM. **I.** Model of KIF11 binding to parallel bundles of minus-end-out microtubules, enacting torque on these bundles; limiting microtubule and dendritic growth. Newly dynamic microtubules are dark green.

To determine whether the observed effects were due to constitutive overexpression of KIF11 or if our pharmacological inhibition interpretations were incorrect, we acutely activated pdDronpa-KIF11 with cyan light for 10 minutes and assessed the impact on dendrites and MT dynamics (Figure 6B, Supplementary Movie S11). Acute activation of KIF11 resulted in a significant reduction in the flux of both plus-end-out (anterograde) and minus-end-out (retrograde) MTs in the targeted dendrites (Figure 6B-D). Additionally, this acute activation led to increased MT kinking and looping (Figure 6B,E-G). Notably, this acute activation did not produce the extensive twisted MTs seen with sustained overexpression (Figure 6H). These findings suggest that KIF11 predominantly binds to parallel MTs in mature neurons, exerting torque that limits MT dynamics and contributes to aberrant dendritic arborization (Figure 6I).

## DISCUSSION

Early in brain development, the organization of microtubules (MTs) and actin dynamics determines whether neurites become axons or dendrites. Axons align MTs with their plus-ends oriented away from the soma, while dendrites maintain a mixed MT polarity (Cheng and Poo, 2012; Lanoue and Cooper, 2019; Ricolo et al., 2021; Sakakibara et al., 2013). As dendrites mature, their MTs become more organized into parallel bundles of like-MTs, guided by cytoskeletal organization and adhesion cues, such as Wnt signaling (Lanoue and Cooper, 2019; Wayman et al., 2006). This organization and stability of MTs and actin are crucial for maintaining and modulating dendritic arborization through both activity-independent and activity-dependent mechanisms (Dent, 2020; Urbanska et al., 2008). In mature dendrites, plus-end-out MTs exhibit dynamic behaviors, whereas minus-end-out MTs are stabilized (Baas and Black, 1990; Kleele et al., 2014; Tas et al., 2017; Yau et al., 2016). This orientation and stability are critical for regulating cargo trafficking and neuronal morphology (Baas and Black, 1990; Feng et al., 2019; Muhia et al., 2016; Tas et al., 2017; Thyagarajan et al., 2022).

Our observations reveal that KIF11 knockdown increases retrograde trafficking of EB3, as evidenced by increased flux but reduced speed (Figure 2H-J). This reduced speed suggests that the retrograde EB3 comets may originate from the minus end of plus-end-out MTs. Patronin/CAMPSAP2, a minus-end binding protein, affects minus-end MT growth in Drosophila dendrites (Cao et al., 2020). However, the interaction between KIF11 and CAMPSAP2 remains unclear. The observed reduction in growth rate should be more substantial—at least fourfold slower—to align with in vitro polymerization rates of the minus end (Strothman et al., 2019). Another possibility is that KIF11 may normally prevent minus-end-out MTs from entering axons from the cell body, though this proposed mechanism of MT sliding has not been observed in neurons (Guha et al., 2021). Given that primary dendrites were unaffected by KIF11 inhibition and the increase in minus-end-out MTs occurred more than 50 µm away from the cell body, this explanation seems unlikely (Figure 1G,H; Figure 2N). Additionally, KIF11 might restrict gamma-tubulin or other nucleating factors from entering axons and initiating minus-end-out MT growth. However, KIF11 is found densely decorating distal MAP2-negative projections in our cultures (Supplementary Figure S1), complicating the understanding of its role in this process. Further studies are needed to explore these possibilities.

Our study found that loss-of-function of KIF11 in mature neurons (DIV14 or later) leads to increased dendritic arborization and KIF11 overexpression decreases dendritic arborization (Figure 2E, Figure 4C-E) (Swarnkar et al., 2018). This contrasts with earlier studies showing decreased arborization following KIF11 loss-of-function in immature neurons (DIV7 or earlier) (Freixo et al., 2018; Kahn et al., 2015). However, others have also found KIF11 overexpression reduces dendritic complexity in mature neurons (Lucero et al., 2022) We attribute this discrepancy to differences in MT organization between maturing and established arbors. In both cases, dendrites exhibit mixed polarity, but in mature neurons, MTs are organized into parallel bundles (Tas et al., 2017). Early in development, KIF11 likely binds to antiparallel MTs, promoting neurite sliding and outgrowth, similar to its role at the mitotic spindle (Ferenz et al., 2010). In mature neurons, KIF11 interacts with parallel MTs, where it cannot promote sliding but may induce rotational forces (Meißner et al., 2024). Given that stable, acetylated MTs are positioned centrally in mature dendrites, the growth from minus-end-out MTs could be central to creating new branches in the absence of KIF11 (Katrukha et al., 2021). It remains unclear whether KIF11 directly organizes parallel MT bundles or if its effects are secondary to other molecular organizers.

Overexpression of KIF11 results in curled MTs, similar to those seen in vitro with a KIF11-KHC chimera lacking the KIF11’s tail domain (Chen and Hancock, 2015). KIF11 was previously proposed to act as an MT polymerase. However, we find it unlikely that KIF11 functions as a polymerase in mature hippocampal neurons. Expression of KIF11^ΔCterm^, WT, and KIF11^Y81F^ all resulted in decreased dynamics of plus- and minus-end-out MTs, rather than an increase. While some observed loops might be due to polymerase activity, and we can’t resolve protofilaments, the compression, and twisting behaviors suggest KIF11 also crosslinks and exerts rotational forces on MTs. Both pharmacological and genetic depletion of KIF11 increase MT dynamics and growth, indicating that KIF11 limits rather than enhances polymerization.

Our study also explored how KIF11 mutations, predicted to cause loss of function, relate to intellectual disabilities. We focused on mutations from patients with minor microcephaly (−2 SD below the mean) yet diagnosed with severe intellectual disabilities, to isolate the effects of KIF11 mutations from those due to microcephaly alone (Jones et al., 2014). We examined mutations in both the N-(KIF11^Y81F^) and C-terminus (KIF11^ΔCterm^) within mature neurons (Figure 4 and Supplementary Figure S4). Expression of MCLID patient mutations KIF11^ΔCterm^ and KIF11^Y81F^ produced phenotypes consistent with force exertion on MTs, and these mutant KIF11 proteins colocalized with MTs, suggesting they retain MT-binding ability. Analysis of these mutants with an MT gliding assay also supports these, as both KIF11^ΔCterm^ and KIF11^Y82F^ retained the ability to bind and slide MTs (Figure 5D). Although it is unclear if they form functional spindles, these findings imply they may still function in mature or developing neurons, contributing to disease phenotypes.

KIF11^Y81F^ expression had nearly indistinguishable effects from WT KIF11 on dendritic arborization, suggesting it is nearly functional. However, in the MT sliding assay KIF11^Y82F^ slowed KIF11 velocity, which would be expected for a rigor mutant (Figure 5D). While Y82F is a conservative mutation, its position near the nucleotide-binding site and tail-binding site may imply that the hydroxyl group is enough to impair the dissociation of the tail, keeping it bound to the microtubule longer (Bodrug et al., 2020). KIF11^ΔCterm^ expression had less impact on dendritic arborization and plus-end-out MT dynamics and showed more diffuse behavior, allowing some visualization of dendritic spines (Supplementary Figure S1H). This indicates that KIF11^ΔCterm^ is a more severe mutation, as the tail domain is essential for coordinating with the motor domain in the homotetramer (Bodrug et al., 2020). Consistent with this, the KIF11^ΔCterm^ displayed an increased velocity in the microtubule sliding assay, similar to the Eg5-Δtail-GFP construct previously studied (Bodrug et al., 2020). The similarity of the velocity between our KIF11^ΔCterm^ (34.2±1.6 nm/s) and the Eg5-Δtail-GFP (33±4 nm/s) also indicates that the deletion of the portion of the BASS domain in the KIF11^ΔCterm^ may not have affected tetramer formation, though this remains to be directly examined. Nevertheless, both mutations appear to retain the ability to exert force on MTs, likely contributing to MCLID pathophysiology.

In summary, we demonstrate that the mitotic kinesin KIF11 binds to parallel MT bundles in mature neurons to regulate minus-end-out MT growth and dendritic branching. This regulation extends beyond dendrites to mature axons. Disruption of MT stability affects synaptophysin, mitochondrial, and lysosomal trafficking in dendrites. Overexpression studies reveal that KIF11 exerts rotational forces in mature neurons. Additionally, patient mutations associated with intellectual disabilities retain the ability to generate force on MTs. Finally, the creation of a photo-inhibitable KIF11 reveals that localized, temporal inhibition is sufficient to increase MT dynamics and dendritic growth. These findings establish KIF11 as MT dynamics rheostat in mature neurons and support a new model for dendritic arborization where minus-end-out MTs initiate new branch points. Future studies should investigate the origin of these dynamic minus-end-out MTs, KIF11’s regulatory partners, and the effects of these changes on neuronal circuitry and behavior in vivo.

## MATERIALS AND METHODS

### Neuronal Cultures and Transfection

For this study, *in vitro* experiments were performed in primary hippocampal cell cultures obtained from CD1 pups. CD1 pregnant females were purchased from Charles River. The experiments were carried out during the light part of the cycle. Housing and experimental procedures were approved and supervised by the Institutional Animal Care and Use Committee of the Wertheim UF Scripps Institute following their Guide for the Care and Use of Laboratory Animals.

Primary hippocampal mouse neuronal cultures were prepared from the brains of embryonic day 16-17 CD1 mice of both sexes. For imaging, cells were plated at 1.7 ×10^5^ on poly-D-lysine-coated (500 μg/ml in Borate Buffer) Mattek dishes. Cultures were plated in Neurobasal medium (Invitrogen) supplemented with 10% fetal bovine serum and penicillin/streptomycin mix and grown in Neurobasal medium supplemented with 2% B27 (Invitrogen), 0.5 mM glutamax, and penicillin/streptomycin mix at 37 °C in 5% CO_2_.

### Fluorescence In Situ Hybridization (FISH)

Two 200– to 300–base pair (bp) fragments of KIF11 cDNA were subcloned into PCRII TOPO vectors and in vitro–transcribed into digoxygenin (DIG)-labeled probes for FISH.. One of the fragments, targeting the 5′ end of KIF11, was more efficient at producing a robust signal and therefore was used for all subsequent studies. DIV17 primary hippocampal cultures (in 35mm Mattek dish) were washed with D-PBS and then fixed in a freshly prepared solution of 4% paraformaldehyde for 10 min. Dishes were then washed with 0.2% glycine in D-PBS (5 min), D-PBS washes (2 x 5 min), acetylated in TEA solution for 10 min, pre-hybridized at 68°C for 1 hr and hybridized with the probes overnight (approx. 16 hrs). After hybridization dishes were washed (3 × 10min) with D-PBS. Then the immunohistochemistry protocol was followed (see below). Antibodies to Map2 (1:1000, Synaptic Systems, 188004) were used to identify dendrites and anti-DIG to recognize probes (1:1000, Roche, 11207733910). TSA Plus Fluorescein Systems from PerkinElmer (TSA plus Cyanine detection kit, Akoya) was used according to the manufacturer’s protocol for DIG detection. Images were acquired using Zeiss ELYRA PS.1 Structured Illumination Super-Resolution Microscopy.

### Immunocytochemistry(ICC) analysis

DIV22 primary hippocampal cultures (in 35mm Mattek dish) were washed with PBS and then fixed in a freshly prepared solution of 4% paraformaldehyde for 15 min. After three or more rinses in PBS, the cells were then incubated in 10% normal horse serum(NHS) (Gibco) in PBS with 0.1% Triton X-100 for 1hr at room temperature to reduce the non-specific binding of primary antibody. Next, the cells were incubated with primary antibodies in 0.1% Triton X-100 1: anti-Kif11 (1:1000, #ab51976, Abcam), anti-Synaptophysin (1:1000, #ab32594, Abcam), anti-Map2 (1:1000, #188004, Synaptic systems),) and anti-alpha Tubulin (1:1000, PA1-38814,ThermoFisher Scientific) overnight at 4°C. Cells were then washed thoroughly with three rinses in PBS followed by a 1 hr incubation in secondary antibodies: Alexa 405/488/568-conjugated secondary antibodies (1:1000; Molecular Probes) for 1 hr. at room temperature. After washing three times with PBS, images were acquired using Zeiss ELYRA PS.1 Structured Illumination Super-Resolution Microscopy.

### Structured Illumination Super-Resolution Microscopy (SIM)

Following immunocytochemistry, neurons were imaged using a Zeiss ELYRA PS.1 instrument (Carl Zeiss, Jena, Germany) at a resolution 1028X1028 pixels, using a Zeiss 63X/1.4 NA Plan Apochromatic objective. Each fluorescent channel, 405, 488 and 561 were acquired using three pattern rotations with 3 translational shifts. The final SIM projection images were reconstructed using Zen 2013 (Carl Zeiss, Jena, Germany) and analyzed using ImageJ.

### EB3 Imaging and Analysis

DIV14-16 Primary hippocampal mouse neurons were simultaneously transfected via combiMag and Lipofectamine with EB3-miRFP703 (Addgene #79994) and either NC-GFP (Origene TR30013) or KIF11-shRNA-C (Origene TG501174). After 24-48hrs of transfection, EB3-miRFP703 imaging was performed at 37°C with 5% CO_2_ in TOKAI HIT STX Stage Top Incubation Chamber, using a confocal microscope (FV3000; Olympus; UPLAPO60XOHR Objective in immersion oil). Time-lapse imaging was acquired with OlyVIA 16-bit Software (Olympus). Pyramidal neurons expressing both EB3-miRFP703 and either NC-GFP of KIF11-ShRNA were selected for imaging. Time-lapse movies were recorded with a zero second delay for _ secs, with 3-frame averaging, first in the far-red channel, then in the green channel.

### EB1 imaging and drug treatment

Primary hippocampal neurons, plated at low density (100K cells in a 35mm Mattek glass bottom dish) are transfected with Dendra2-EB1 (Addgene Plasmid #57714) via lipofectamine DIV15-17. Six to twelve hours after transfection feeding media is replaced with pre-warmed Hibernate-E. Immediately after replacing the media, the dish is placed in the prewarmed Tokai Hit stage top incubator at 37°C. The cells are then imaged with the Olympus Fluoview FV3000 confocal microscope with the cellSens V4.1 software (64-bit) with the UPlanApo 60x/1.5 Oil N.A. objective, and with a 488 nm (20 mW) laser. Healthy cells that express Dendra-EB1 that have a 50µm region of axon or dendrite in the same focal plane are selected. These neurons are then recorded in the 488 channel for 2 minutes at 1.25 frames per second. These movies are then opened in FIJI ImageJ and the ROI is manually drawn along the in-focus axon/dendrite and processed with the KymoResliceWide plugin. The slope of the EB3 tracks seen on the kymograph, along with the directionality (moving towards(retrograde) or away(anterograde) from the soma), the length of the tracks, and the number of tracks are manually measured with the measure tool in FIJI ImageJ. These values were then used to calculate the MT growth rate (µm/sec), the orientation of the MT (anterograde track= plus-end-out, retrograde tracks=minus-end-out), the duration of the growth, and the frequency of the growth, respectively.

### Sholl Analysis

After 72 hours of transfection of hippocampal neurons using shRNA plasmid expressing eGFP, images of dendrites were collected at room temperature in the light microscopy facility at the Max Planck Florida Institute, using a confocal microscope (LSM 780; Carl Zeiss; Plan Neofluor 63X/1.3 N.A. Korr differential interference contrast M27 objective in water). Z-stack images were acquired using ZEN 2015 (64-bit) software (Carl Zeiss) and dendritic arbors were manually traced using confocal projection images and later quantified by Sholl analysis FIJI (ImageJ, NIH). The center of the soma is considered as the midpoint and the origin of the concentric radii was set from that point to the longest axis of the soma. The parameters set for analysis were: starting radius 20 μm, ending radius 100 μm, radius step size 10 μm. The maximum value of sampled intersections reflecting the highest number of processes/branches in the arbor was calculated and the number of intersections was plotted against the distance from the soma center in μm. Data was analyzed using one-way ANOVA with Tukey post hoc test.

### Mitochondrial and lysosomal vesicle imaging and drug treatment

Primary hippocampal neurons, plated at low density (100K cells in a 35mm Mattek glass bottom dish) and imaged at DIV16. Prior to imaging, feeding media is replaced with pre-warmed Hibernate-E containing 50nM MitoTracker Green and 50nM LysoTracker Red. After a 30-minute incubation, the media is then replaced with pre-warmed Hibernate-E and either DMSO or 1µM Ispinesib or BRD9876. After a 15-minute incubation, the dish is placed in the prewarmed Tokai Hit stage top incubator at 37°C. The cells are then imaged with the Olympus Fluoview FV3000 confocal microscope with the cellSens V4.1 software (64-bit) with the UPlanApo 60x/1.5 Oil N.A. objective and with a 488 nm (20 mW) and 561 nm: 20mW) laser. Healthy cells that contained clearly stained mitochondria and lysosomal vesicles and a 100µm region of axon or dendrite in the same focal plane are selected. These neurons are then recorded first in the 568 channel and then in the 488 channel for 2 minutes at 1.25 frames per second. These movies are then opened in FIJI ImageJ and the ROI is manually drawn along the in-focus axon/dendrite and processed with the KymoResliceWide plugin. The number and directionality (moving towards(retrograde) or away(anterograde) from the soma of tracks per 100µm to determine flux.

### Optical manipulation of KIF11

The pdDronpaV-KIF11 construct was custom synthesized with mouse Kif11 (NM_010615.1) with N-pdDronpaV tag (Zhou et al., 2017) with a custom linker (GSSGGGGSGGGGSGGGGS), in the lentiviral vector pEZ-Lv236 with CAG promoter. The pdDronpa-CDK5 (pLL3.7m-psCDK5, Addgene #89365) construct was used as a control for optimizing pdDronpa association and dissociation.

HEK 293T cells were maintained in high glucose Dulbecco’s Modified Eagle Medium (DMEM, Thermo Fisher) supplemented with 5% fetal bovine serum (FBS, Thermo Fisher) and 2 mM glutamine (Sigma, Thermo Fisher) at 37°C in air with 5% carbon dioxide. HEK 293T cells were transfected at 75-90% confluency with Lipofectamine 2000 (Invitrogen) in 35mm 1.5 coverglass bottom dishes MatTek dishes. Primary Hippocampal neurons were plated and transfected as previously described for EB3 imaging.

Cells were imaged within 16 hours of transfection using an Olympus Fluoview FV3000 confocal microscope with the cellSens V4.1 software (64-bit) with the UPlanApo 60x/1.5 Oil N.A. objective. The imaging protocol for pdDronpa monomerization and dimerization was first optimized in HEK293T cells starting with the protocol outlined in (Zhou Xin et al., 2017) and adjusted to prevent cell death, minimal photobleaching, and a clear increase in fluorescence intensity with 405nm dimerization. pdDronpaV was imaged with a 10% neutral density filter, the 485/30-nm excitation filter, the 505-nm dichroic mirror, and the 525/40-nm emission filter. miRFP703-EB3 was imaged with a 10% neutral density filter, a 640-nm excitation filter, a 730-nm dichroic mirror, and a 640/730-nm emission filter. Dronpa variants were photoswitched off (enabling KIF11 activation) using the same excitation and dichroic filters as for imaging but without a neutral density filter for 5min. DronpaV was reactivated (enabling KIF11 inactivation) with 405nm (50 mW) at 0.1% power for 5min.

Fluorescence was analyzed in ImageJ by measuring corrected total cell fluorescence (CTCF) = Integrated Density – (Area of selected cell X Mean fluorescence of background readings) and normalizing values to Frame 0sec. The intensities and durations of light used in each experiment were noted in the figures. EB3 Flux was calculated as described above. New branch points were characterized from maximum projection intensity images from the FarRed (EB3) channel, as new growth longer than 5µm. Loop number and size was also calculated from maximum projection intensity images from the FarRed (EB3) channel.

### Microtubule sliding assay with KIF11 mutants: Gene expression and protein purification

KIF11 expression and purification was conducted based on previous work (Meißner et al., 2024). Viruses were generated using the FlexiBac system (Lemaitre et al., 2019). SF9 cells (IPLB Sf21-AE, Merck 71104) at 1 million cells per mL were infected with virus (1:100, v/v) and genes were expressed for 96 h at 27 °C and 120 rpm. Cells were centrifuged with 300 × g for 10 min at 4 °C. Pellets were resuspended in PBS (1% of expression volume) with protease inhibitor, flash frozen in liquid nitrogen, and stored at −80 °C. For purification, cell pellets were thawed on ice and resuspended in purification buffer (50 mM NaH2PO4, 300 mM KCl, 2 mM MgCl2, 0.5 mM TCEP, 0.5 mM ATP, pH 7.5) with protease inhibitor. The lysate was cleared with an ultracentrifuge spin with 40,000 rpm for 1 h at 4 °C. The supernatant was filtered through a 0.45 µm filter and loaded on a 1 mL HiTrap column systempump. The column was washed with immobilized metal affinity chromatography (IMAC) wash buffer (purification buffer with 20 mM imidazole) and the protein was eluted with IMAC elution buffer (purification buffer with 300 mM imidazole) with an elution gradient. Protein-containing fractions were pooled and concentrated with Amicon filters (cutoff 100 kDa). 3C protease was added (1:100, v/v) and the His 6 tag was cleaved overnight at 4 °C. The protein solution was diluted 6-fold to reduce the imidazole concentration and passed over the HiTrap column again. The flow through was concentrated to 0.5 mL, cleared at 17,000 × g for 10 min, and gel filtered over a Superose6 column with purification buffer. 5% glycerol was added and the protein was flash-frozen in liquid nitrogen and stored at −80 °C.

### Microtubule sliding assay with KIF11 mutants: 2D sliding experiments

To assemble 6 flow chambers, 7 strips of Nescofilm were placed on the coverslip with structures and covered with a regular coverslip with DDS coating. The Nescofilm was melted on a hot plate. Every second channel was sealed with two component silicone (Twinsil picodent) to avoid cross-contamination of channels. Channels were flushed with: (I) 0.2 mg/mL TAMRA 5G5 antibody (Invitrogen, RRID AB_2536728) solution in PBS for 5 min, (II) 1% F127 (w/v in PBS) solution for at least 1 h, (III) 3 washes with BRB80, (IV) fixed microtubules in motility buffer (MB-ADP, BRB80 with 10 µM taxol, 200 µg/mL casein (from bovine milk, Sigma C7078), 10 mM DTT, 0.1% (v/v) Tween-20, 20 mM D-glucose, 1 mM ADP), (V) MB-ADP wash, (VI) KIF11 (10nM KIF11-EGFP; 80nM KIF11ΔCterm-mNG or 15nM KIF11Y82F-eGFP) in MB-ADP for 5 min, (VII) MB-ADP wash, (VIII) cargo microtubules in MB-ADP, (IX) wash with MB-ADP + + (MB-ADP with 200 µg/mL glucose oxidase and 20 µg mL−1 catalase), (xii) MB-ATP + + (MB-ADP + + with 1 mM ATP instead of ADP).

### Microtubule polymerization

Microtubule polymerization Tubulin was purified from pig brains according to standard protocols (Castoldi and Popov, 2003) and labeled in house. Briefly, tubulin was polymerized and microtubules were isolated by centrifugation in a glycerol cushion. Microtubules were mixed with the labeling dye (TAMRA succinimidyl ester, ThermoFischer; Alexa Fluor 488 Carboxylic Acid, 2,3,5,6-Tetrafluorophenyl ester, ThermoFischer; ATTO 647N Amine-reactive NHS-ester, ATTO-TEC) in a 10 to 20-fold molar excess of the dye and incubated for 30 min. The reaction was quenched with 0.5 M potassium glutamate and the tubulin was purified with depolymerization-polymerization cycles. Labeled tubulin was mixed in a ratio of 1:3 with unlabeled tubulin for experiments. Fixed and cargo microtubules were grown with tubulin with guanylyl-(α,β)-methylene-diphosphonate (GMP-CPP) and stabilized with taxol. For fixed microtubules, 40 µL elongation mix containing 1.25 mM GMP-CPP, 1.25 mM MgCl 2 and 4.5 µM TAMRA labeled tubulin in BRB80 (80 mM Pipes at pH 6.9, 1 mM MgCl 2, 1 mM EGTA) was incubated on ice for 5 min and then for 30 min at 37 °C. Microtubules were pelleted (17,000 × g, 15 min) and resuspended in elongation mix (1.25 mM GMP-CPP, mM MgCl 2 and 0.5 µM TAMRA labeled tubulin in BRB80) and grown for 2–3 days at 32 °C. Microtubules were pelleted (17,000 × g, 8 min) and gently resuspended in BRB80 with 10 µM taxol (BRB80X). They were kept at room temperature for several days for annealing. To grow cargo microtubules, a polymerization mix with 1.25 mM GMP-CPP, 1.25 mM MgCl 2 and 5 µM Atto647N labeled tubulin was incubated on ice for 5 min and then for 8 min at 37 °C. Microtubules were pelleted (17,000 × g, 15 min) and resuspended in BRB80X.

### Optical image acquisition

Optical image acquisition Optical imaging was performed using an inverted fluorescence microscope (Axio Observer Z1; Carl Zeiss Microscopy GmbH) with a 63× oil immersion 1.46NA objective (Zeiss) in combination with an EMCDD camera (iXon Ultra; Andor Technology) controlled by Metamorph (Molecular Devices Corporation). A LED white light lamp (Sola Light Engine; Lumencor) in combination with a TRITC filterset (ex 520/35, em 585/40, dc 532: all Chroma Technology Corp.), an Atto647N filterset (ex 628/40, em 692/40, dc 635) and a GFP filter set (ex 475/35, em 525/45), corresponding to TAMRA labeled microtubules, Atto647N labeled microtubules and EGFP labeled motors/Atto488 labeled microtubule extensions, respectively, were used for epifluorescence imaging. The sliding of cargo microtubules was imaged in the Atto647N channel for 10 min with 10 frames/s with an exposure time of 100 ms. Fixed microtubules were imaged in the TRITC channel for 100 frames at 10 frames/s with an exposure time of 100 ms after imaging the cargo microtubules.

### Statistical analysis

Levels of significance in this study are based on *p*-values calculated by GraphPad Prism 9 (Graph Pad Software) and derived using Student’s t-test, Mann-Whitney U test, One-way or Two-way ANOVA. Tukey, Dunnett, or Sidak tests were used for post hoc analyses. Significance was defined as < 0.05. Outliers were defined with a Grubb’s test (Alpha = 0.05). The details for each experiment and the number of replicates and statistical specifications are indicated in the figure legends, results, and Supplementary Data.

## Acknowledgments

We gratefully acknowledge this work was supported by NIH grants (5R01MH094607-05, 1F32MH131420-01A1, and 1R01MH119541-01A1, R01MH118444), funding support from NSF (Award Number: 2231247) and a Training Grant in Alzheimer’s Drug Discovery from the Lottie French Lewis Fund of the Community Foundation for Palm Beach and Martin Counties, Florida. We sincerely thank Ms. Emelie Lewis for her help with ICC experiments, and Drs. HaJeung Park and Kevin Knight for modeling the KIF11 structure, and Dr. Laura Meißner for initial discussions of microtubule sliding experiments.

## Data availability

All data supporting the findings of this study are available within this manuscript and its Supplementary Information.

## Author Contributions

J.W., and S.P. conceived of and designed experiments. J.W. conducted all the experiments and analyzed the data except the MT sliding assays. LN conducted MT sliding and KIF11 motility measurements under the supervision of SD. J.W wrote and revised the manuscript with inputs from all the authors. SP supervised the study. All authors read and commented on the manuscript.

