## Supplementary Files for "Intellectual disability-causing mutations in KIF11 impair microtubule dynamics and dendritic arborization"

[*]Corresponding author:

4 Supplementary Figures (S1-4)
11 Supplementary Movies (S1-11)
Link for Movies
https://www.dropbox.com/scl/fo/1si5qwbjqs9cdfobb6fww/ABrXTVendetBBTfJv3xLdAU?rlkey=6uqjysrkn9v7e58q89f9dorjw&st=6bi8g1bj&dl=0

### SUPPLEMENTARY FIGURES

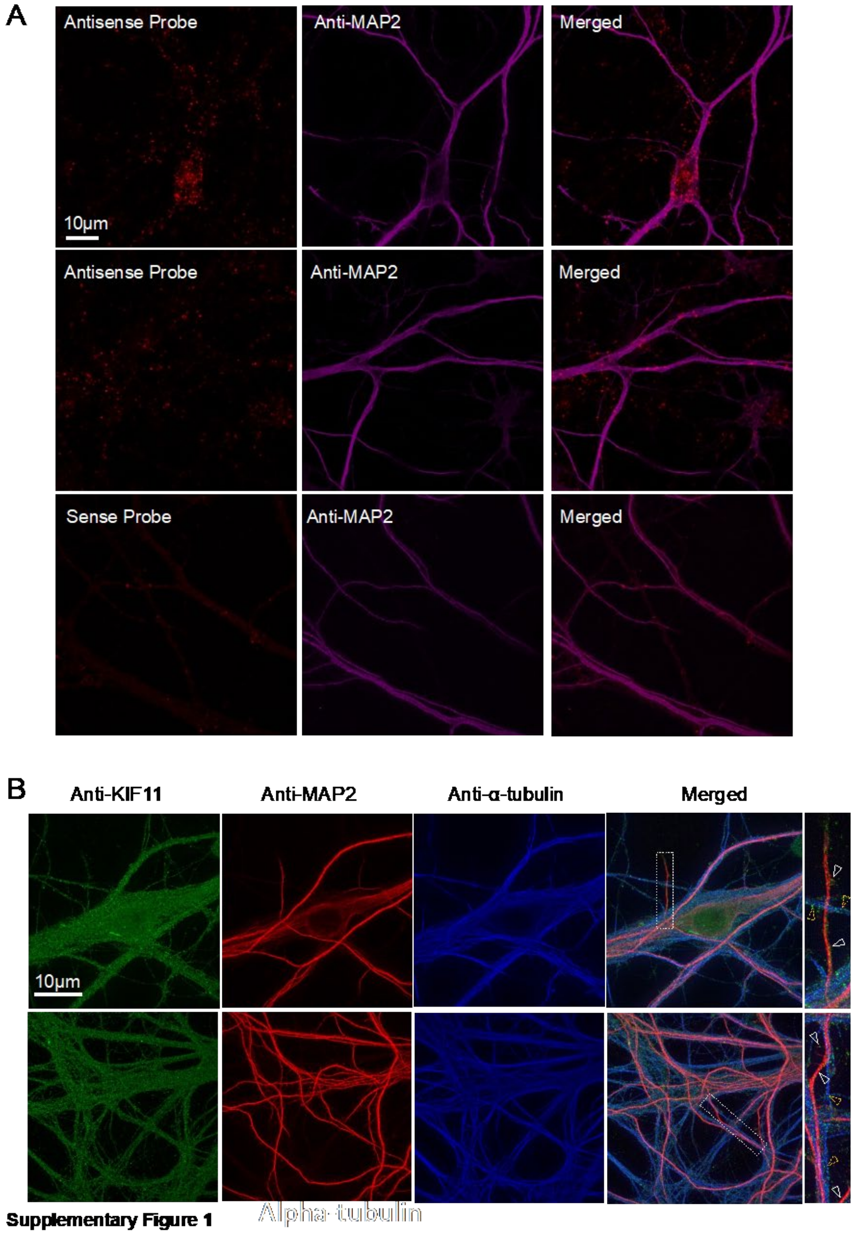


**Supplementary Figure S1. Localization of KIF11 mRNA and protein in mature neurons. A.** Fluorescence In Situ Hybridization (FISH) images of DIV17 primary hippocampal neurons. Antisense probe shows KIF11 mRNA to colocalize with MAP2 staining in the cell body(top) and distal dendrites(middle). Sense probe shows comparatively less staining, used as a control. KIF11(red) and dendritic marker MAP2(magenta). Scale bars = 10 µm. **B.** Immunocytochemistry (ICC) images of DIV22 primary hippocampal neurons probing for KIF11(green) MAP2(red) and alpha-tubulin(blue). Inset images show KIF11 to localize to both MAP2-positive(white arrowheads) and MAP2-negative(yellow arrowheads) projections both proximally (top) and distally(bottom) from the cell body.

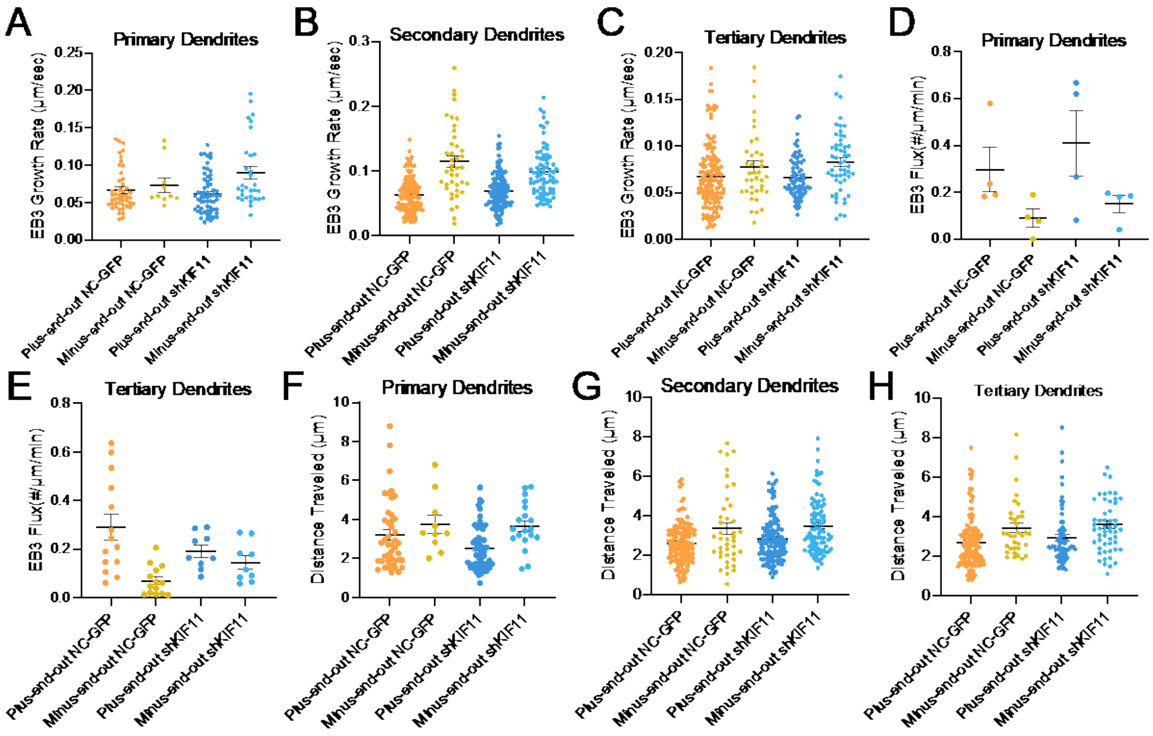


**Supplementary Figure S2. Analysis of KIF11's impacts on microtubule dynamics per branch type. A-H** Segmented analysis of data set described in Figure 2**. A, B, C**. EB3-comet growth rate for plus-end-out and minus-end-out EB3 comets in primary (**A**), secondary (**B**), and tertiary dendrites (**D**). One-way ANOVA, Tukey's multiple comparison test. Error bars=SEM. **E, F, G**. EB3-comet flux for plus-end-out and minus-end-out EB3 comets in primary (**E**), secondary (**F**), and tertiary dendrites (**G**). One-way ANOVA, Tukey's multiple comparison test, *p-value<0.05. Error bars=SEM. **H, I, J**. EB3-comet growth distance for plus-end-out and minus-end-out EB3 comets in primary (**H**), secondary (**I**), and tertiary dendrites (**J**). One-way ANOVA, Tukey's multiple comparison test. Error bars=SEM

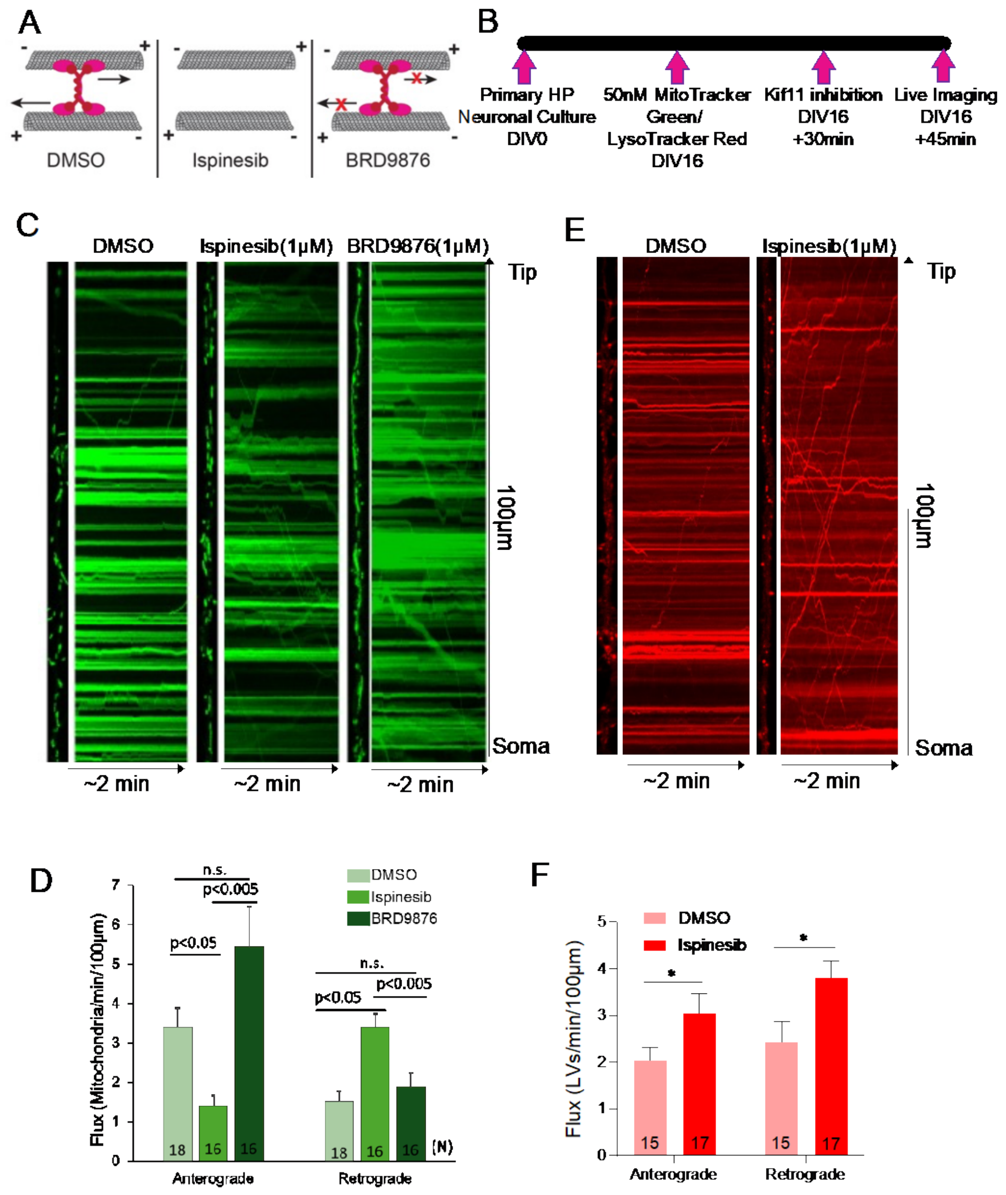


**Supplementary Figure S3. Transient KIF11 inhibition disrupts the transport of both mitochondria and lysosomal-related organelles.**

**A.** Experimental timeline. **B**. Schema of how two different KIF11 inhibitors impact KIF11's ability to bind microtubules. Ispinesib prevents KIF11 from binding microtubules. BRD9876 locks KIF11 on microtubules bound to the microtubule. **C.** Selected dendrites and their corresponding kymographs from neurons stained with MitoTracker Green. **D**. Mitochondria flux from C for both anterograde and retrograde tracks. One-way ANOVA, Tukey's multiple comparison test, n.s.= not significant, Error bars=SEM. **E.** Selected dendrites and their corresponding kymographs from neurons stained with Lysotracker Red. **F.** Lysosomal Related Organelle (LRO) Flux from E for both anterograde and retrograde tracks. One-way ANOVA, Tukey's multiple comparison test, **p-value <0.05, Error bars=SEM. N=Number of dendrites analyzed.

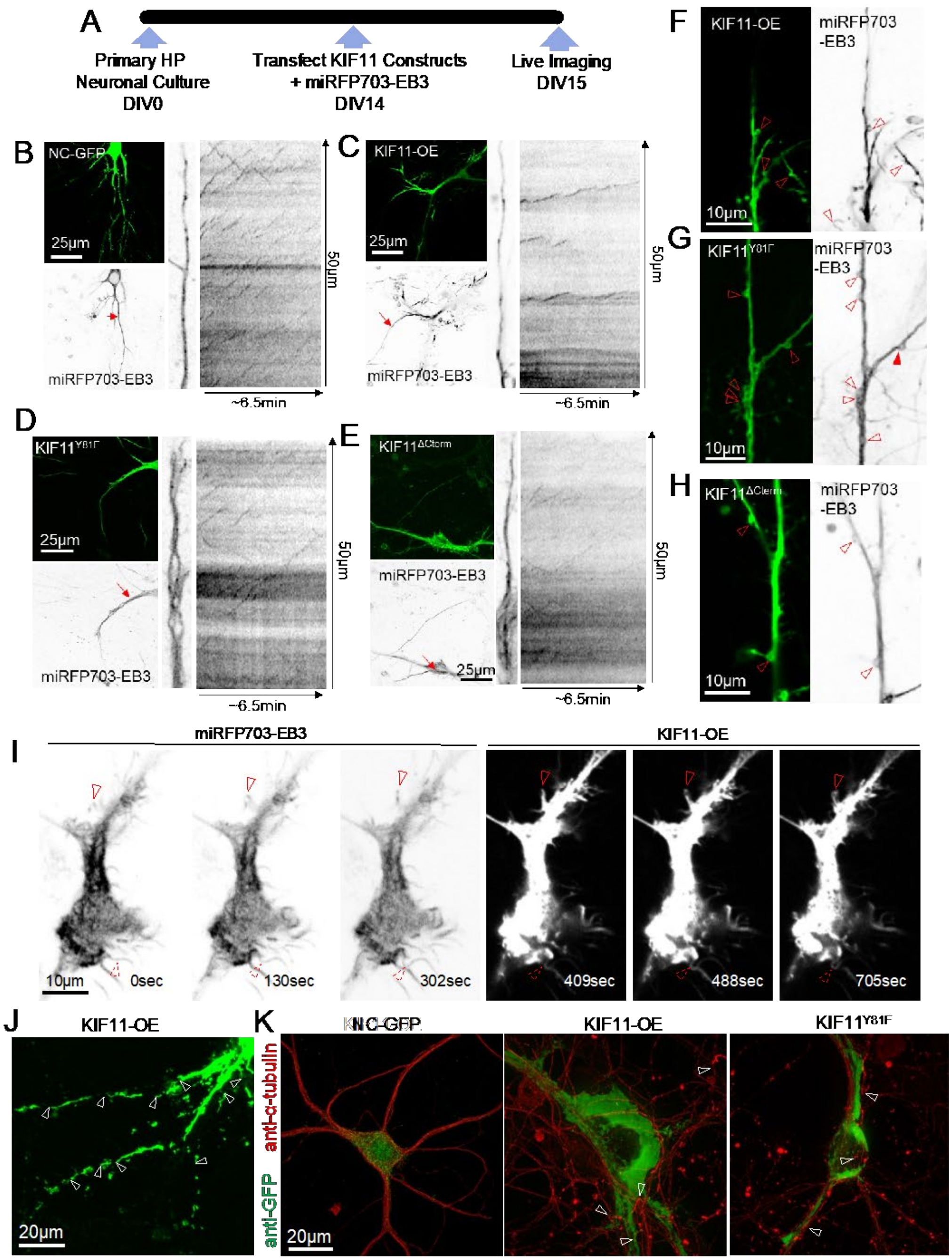


**Supplementary Figure S4. KIF11 patient mutations microtubule dynamics.**

**A.** Experimental timeline**. B-E.** Representative Kymographs for Figure 3F-L. **B.** Left: Maximum projection intensity images of a dendrite from neurons co-expressing NC-GFP (scrambled negative control) and miRFP703-EB3. Right: Selected dendrite and kymograph of miRFP703-EB3. **C.** Left: Maximum projection intensity images of a dendrite from a neuron co-expressing KIF11-OE and miRFP703-EB3. Right: Selected dendrite and kymograph of miRFP703-EB3. **D.** Left: Maximum projection intensity images of a dendrite from a neuron co-expressing KIF11$^{Y81F}$

and miRFP703-EB3. Right: Selected dendrite and kymograph of miRFP703-EB3. **E.** Left: Maximum projection intensity images of a dendrite from a neuron co-expressing KIF11$^{\Delta Cterm}$ and miRFP703-EB3. Right: Selected dendrite and kymograph of miRFP703-EB3. **F.** Maximum projection intensity images of a dendrite from a neuron co-expressing KIF11-OE and miRFP703-EB3. Empty arrowheads identify loops without EB3 comets. Filled arrowheads identify loops with EB3 comets. **G.** Maximum projection intensity images of a neuron co-expressing KIF11$^{Y81F}$ and miRFP703-EB3. Empty arrowheads identify loops without EB3 comets. Filled arrowheads identify loops with EB3 comets. **H.** Maximum projection intensity images of a dendrite from a neuron co-expressing KIF11$^{\Delta Cterm}$ and miRFP703-EB3. Empty arrowheads identify loops without EB3 comets. Filled arrowheads identify loops with EB3 comets. **I.** Individual frames of a movie of a neuron co-expressing KIF11-OE and miRFP703-EB3. The first ~6.5 minutes of this movie were captured in the far-red channel (miRFP703-EB3) and the next ~6.5 minutes were captured in the green channel (KIF11-OE). An upper empty arrowhead identified a bent neuronal process twisting around itself to form a complete loop within 11 minutes of imaging (twisted loop). The lower dashed arrowhead identifies a neuronal process that starts almost completely straight but bends and nearly completes loop formation within 11 minutes of imaging. **J.** Maximum projection intensity images of a dead neuron expressing KIF11-OE. Arrowheads point to broken and fragmented dendrites. **K.** ICC images of DIV16 primary hippocampal neurons expressing NC-GFP, KIF11-OE, or KIF11$^{Y81F}$(green) and probed for alpha-tubulin(red). White arrowheads point to fragmented and twisted microtubules.

#### SUPPLEMENTARY MOVIES

**File Name: Movie S1**

Description: Representative time-lapse movies of DIV16 primary hippocampal neurons transfected with Dendra-EB1 treated with DMSO(top), Ispinesib(middle), or BRD9876(bottom). Playback speed 60x real time. Total dendrite length=100µm. Related to Figure 1D.

**File Name: Movie S2**

Description: Representative time-lapse movies of DIV16-17 primary hippocampal neurons expressing miRFP703-EB3 and NC-GFP or shKIF11. Playback speed 30x real time. Total length=50µm. Related to Figures2B-E.

**File Name: Movie S3**

Description: Representative time-lapse movies of DIV16-17 primary hippocampal neurons expressing mRuby-Synaptophysin and NC-GFP or shKIF11. Playback speed 30x real time. Total dendrite length=50µm. Related to Figures3B-C.

**File Name: Movie S4**

Description: Representative time-lapse movies of DIV16-17 primary hippocampal neurons expressing mApple-PSD95 and NC-GFP or shKIF11. Playback speed 30x real time. Total dendrite length=50µm. Related to Figures3I-J.

**File Name: Movie S5**

Description: Representative time-lapse movies of DIV16 primary hippocampal neurons stained with MitoTracker Green and treated with DMSO(top), Ispinesib(middle), or BRD9876(bottom). Playback speed 30x real time. Total dendrite length=100μm. Related to Supplementary Figure S3C.

**File Name: Movie S6**

Description: Representative time-lapse movies of for DIV16 primary hippocampal neurons stained with LysoTracker Red and treated with DMSO(top) or Ispinesib(bottom). Playback speed 30x real time. Total dendrite length=100μm. Related to Supplementary Figure S3D.

**File Name: Movie S7**

Description: Representative time-lapse movies of DIV15 primary hippocampal neurons miRFP703-EB3 and NC-GFP, KIF11-OE, $KIF11^{Y81F}$, or $KIF11^{\Delta Cterm}$. Playback speed 30x real time. Total dendrite length=50μm. Related to Supplementary Figure S4B-E.

**File Name: Movie S8**

Description: Representative time-lapse movies of a DIV16 primary hippocampal neuron expressing miRFP703-EB3 and KIF11-OE. Playback speed 30x real time. Total movie width=60μm. Related to Figure Supplementary S4I.

**File Name: Movie S9**

Description: Representative time-lapse movies of HEK293T cells expressing miRFP703-EB3 and pdDronpa-CDK5 or pdDronpaV-KIF11. Playback speed 30x real time. Total movie width=90μm. Related to Figure 5B.

**File Name: Movie S10**

Description: Representative time-lapse movies of a primary hippocampal neuron expressing miRFP703-EB3 and pdDronpaV-KIF11 with 405nm stimulation to inactivate KIF11. Playback speed 30x real time. Total movie width=80μm. Related to Figure 6D.

**File Name: Movie S11**

Description: Representative time-lapse movies of a primary hippocampal neuron expressing miRFP703-EB3 and pdDronpaV-KIF11 with 488nm stimulation to activate KIF11. Playback speed 30x real time. Total movie width=80μm. Related to Figure 7B.
